# NAToRA, a relatedness-pruning method to minimize the loss of dataset size in genetic and omics analyses

**DOI:** 10.1101/2021.10.21.465343

**Authors:** Thiago Peixoto Leal, Vinicius C Furlan, Mateus Henrique Gouveia, Julia Maria Saraiva Duarte, Pablo AS Fonseca, Rafael Tou, Marilia de Oliveira Scliar, Gilderlanio Santana de Araujo, Camila Zolini, Maria Gabriela Campolina Diniz Peixoto, Maria Raquel Santos Carvalho, Maria Fernanda Lima-Costa, Robert H Gilman, Eduardo Tarazona-Santos, Maíra Ribeiro Rodrigues

## Abstract

Genetic and omics analyses frequently require independent observations, which is not guaranteed in real datasets. When relatedness cannot be accounted for, solutions involve removing related individuals (or observations) and, consequently, a reduction of available data. We developed a network-based relatedness-pruning method that minimizes dataset reduction while removing unwanted relationships in a dataset. It uses node degree centrality metric to identify highly connected nodes (or individuals) and implements heuristics that approximate the minimal reduction of a dataset to allow its application to large datasets. NAToRA outperformed two popular methodologies (implemented in software PLINK and KING) by showing the best combination of effective relatedness-pruning, removing all relatives while keeping the largest possible number of individuals in all datasets tested and also, with similar or lesser reduction in genetic diversity. NAToRA is freely available, both as a standalone tool that can be easily incorporated as part of a pipeline, and as a graphical web tool that allows visualization of the relatedness networks. NAToRA also accepts a variety of relationship metrics as input, which facilitates its use. We also present a genealogies simulator software used for different tests performed in the manuscript.

---

In omics research, we frequently apply methods that require independent observations. However, when these observations are individuals from a population, they may be relatives (i.e. not independent). A common solution is to exclude all or part of relatives to reduce dependence, but more efficient solutions are needed to reduce dataset pruning (Supplementary Information: Section 1, SI:S1). We present a relatedness-pruning method based on Complex Network Theory called NAToRA (Network Algorithm To Relatedness Analysis), which simultaneously minimizes the number of observations to be excluded from datasets, increasing their independence.

NAToRA is an algorithm that minimizes the number of individuals to be removed from a dataset. In the context of complex network theory, NAToRA finds the maximum clique in the complement networks. However, because this is an NP-Complete Problem [1], and it is computationally infeasible in several cases, we developed an heuristic version of NAToRA that approximates the minimal reduction of the dataset. In general NATORA models relatives as a network in which individuals (or more in general, observations) are nodes and relatedness coefficients between them are weights of their connections (or edges). In this network, genetically-related individuals called network families are sets of nodes that have at least one sequence of edges connecting all of them. Contrarily, unrelated individuals (or related below a cutoff value) are represented by disconnected nodes. The algorithm receives two inputs: (i) an adjacency list containing pairs of individuals and their relatedness coefficients (Figure 1(a), SI:S2), and (ii) a relatedness cutoff value indicating the maximum of the relatedness coefficient to be allowed after pruning (e.g., corresponding to third-degree kinship and below, Table S1). NAToRA creates a network containing only the individuals linked by relatedness coefficients greater than the cutoff value provided by the user (Figure 1(b), illustrating a third-degree cutoff). From this network, the algorithm first detects and reports network families from the matrix of relatedness coefficients (an information that may be used as a categorical variable in different instances). Then, for each detected family, the heuristic algorithm iteratively prunes individuals with more links than others (i.e., with higher node degree centrality (NDC), [1]) (Figure 1(c), (d), (e), (f)). NDC is a node metric based on its number of edges and it was chosen after comparisons with alternative metrics (SI: S3-S4). If there are individuals with equal NDC, NAToRA prunes those with the highest sum of its edges’ weights. If there is another tie, the algorithm removes one of them randomly. The output is a list of individuals to be excluded from the dataset (Figure 1(g)). These comparisons were performed using pseudo-genealogies generated by a *genealogy simulator* that we developed, described in detail in SI:S3. This simulator aims to create genealogy with reproductive behavior similar to expected in human populations based on parameters provided by the user, allowing to create several different scenarios. After generating the genealogy, the algorithm calculates the theoretical kinship coefficient (Table S1) among all pairs of related individuals.

**Figure 1.**
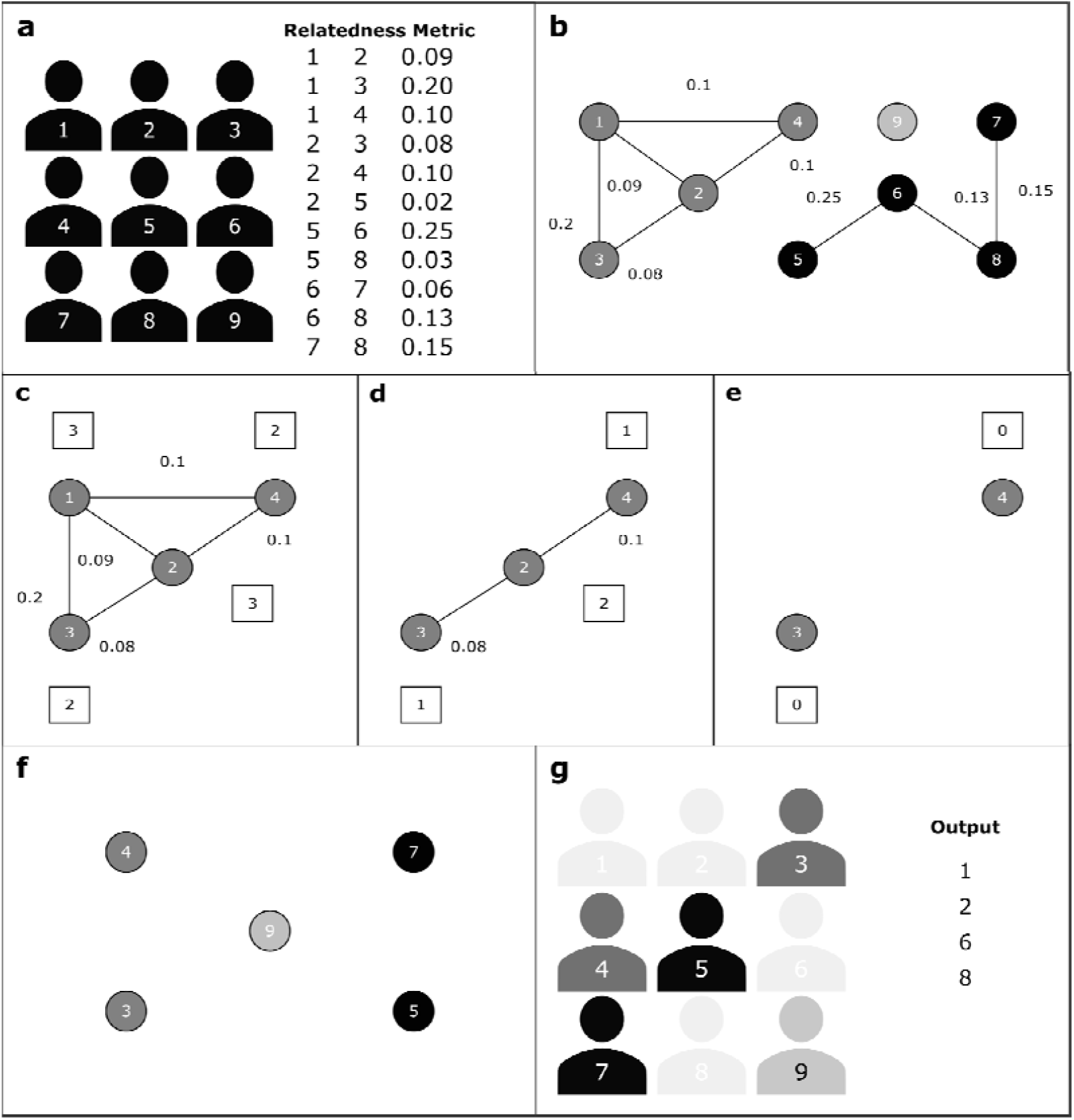
Overview of NAToRA’s (Network Algorithm To Relationship Analysis) network-based algorithm. (a) input file with relatedness metrics for pairs of individuals. (b) relatedness network with kinship cutoff of 0.07; grey-scale colours represent families of genetically-related individuals. (c), (d) and (e) show the node elimination process for the dark grey family component, in which individuals with the highest node centrality degree (NCD, denoted in whit boxes) are iteratively removed (in this case the individuals 1 and 2 with NCD=3). (f) relatedness-pruned network. (g) output file with a list of individuals to be removed from the dataset.

We tested NATORA using relatedness matrices constructed from three genome-wide datasets including related individuals: (i) The Bambuí Cohort Study of Aging (BAMBUÍ) (n=1,442 admixed brazilians) [2], (ii) Matsiguenkas indigenous from the Peruvian Amazon Yunga (SHIMAA) (n=45) [3], (iii) Guzerá *Bos indicus* dairy cattle from the brazilian National Breeding Program (GUZERÁ) (n=1,036) [4] (SI: S5). The study was approved by the Institutional Review Board of the participating institutions.

Overall, NAToRA performs better than relatedness-pruning methods implemented in popular genetics software PLINK v1.90b6.9 [5] and KING 2.2.4 (Kinship-based INference for Genome-wide association studies) [6], showing the best combination of effective relatedness-pruning by removing all unwanted relationships while keeping the largest possible number of individuals in all datasets (Table 1, SI: S7). Specifically, PLINK showed the highest number of remaining individuals in all datasets but it did not exclude all relationships above the relatedness cutoff. KING and NAToRA had similar performances for Bambuí and Guzerá datasets, but NAToRA kept more individuals and KING did not exclude any related individuals in the Shimaa dataset. Also, to assess the impact of pruning individuals to the overall dataset genetic diversity, we analyzed allele frequency patterns and principal components before and after the pruning process. The NAToRA methodology maintains a large part of the variability in all analyzes, showing a better or comparable performance to PLINK and KING, while the latter two softwares do not guarantee the removal of the entire relationship from the dataset (SI: S8).

**Table 1.** Comparison between PLINK, KING and NAToRA relatedness-pruning methods. Values show the percentage of dataset reduction and relatedness reduction. NA values indicate that the method did not work. Bold values indicate the best result in each pairwise comparison. We used 2nd-degree kinship (0.0884) as the cutoff value.

| Metric | PI_HAT |  |  |  | Kinship Coefficient |  |  |  |
| --- | --- | --- | --- | --- | --- | --- | --- | --- |
|  | Sample reduction |  | Relatedness reduction |  | Sample reduction |  | Relatedness reduction |  |
|  | NAToRA | PLINK | NAToRA | PLINK | NAToRA | KING | NAToRA | KING |
| BAMBUÍ<br>N= 1,442 | 39.7% | **37.0%** | **100.0%** | 85.0% | **34.2%** | 41.8% | **100.0%** | 99.9% |
| SHIMAA<br>N=42 | 48.9% | **42.2%** | **100.0%** | 92.0% | **44.4%** | NA | **100.0%** | NA |
| GUZERÁ<br>N=1,036 | 83.0% | **79.7%** | **100.0%** | 98.0% | **79.0%** | 91.5% | **100.0%** | **100.0%** |

NATORA presents three additional advantages. First, its flexibility in accepting different similarity metrics for relatedness-pruning (SI: S6-S7), while PLINK’s and KING’s pruning methods are tied to their own metrics of relatedness (Table 1). For example, NAToRA is also compatible with relatedness metrics calculated by the REAP method (Relatedness Estimation in Admixed Populations) [7], which is more appropriate for admixed populations than PLINK and KING.

Second, although NAToRA provides an alternative to PLINK and KING’s relatedness-pruning methods, it can still be used in pipelines that include broader use of these software, such as genome-wide association testing. For example, one can use PLINK, KING, or other software to perform data quality control and to calculate relatedness metrics, and include NAToRA in the relatedness-pruning step (see NAToRA’s User Guide, SI:S9) to minimize dataset reduction.

Third, NAToRA also provides a function that partitions the dataset in subsets of unrelated individuals, without excluding any individual, for analyses that can be performed with subsets of independent data that can be later combined, as in ADMIXTURE ancestry analysis in [8] (SI: S10). Importantly, other applications of NAToRA rely on its identification of individuals with the highest centrality in a network. These individuals may be conceived as a reduced set of the most representative individuals of their families. This concept, for instance, may be applied in conservation genetics of small natural populations, to select individuals for reproduction. In omics research, this application may allow to select representative individuals or observations for follow-up experiments.

## Conclusions

Considering the importance of the number of individuals (observations) to gain statistical power, NAToRA provides both, a minimal reduction of sample size and an effective removal of undesired kinship relationships. NAToRA is freely available, both as a standalone tool that can be easily incorporated as part of an analysis pipeline, and as a graphical web tool that allows visualization of the relatedness networks.

## Availability of supporting source code and requirements

Project name: NAToRA

Project home page: https://github.com/ldgh/NAToRA_Public

Operating system(s): Platform independent

Programming language: NAToRA was implemented in Python and the the scripts that compose the NAToRA toolkit was implemented in Perl

Other requirements: Python3 or higher and library NetworkX 2.0 or higher

License: GNU

## Data Availability

All the data used on this work is freely available at https://github.com/ldgh/NAToRA_Public on Datasets folder

## Authors’ contributions

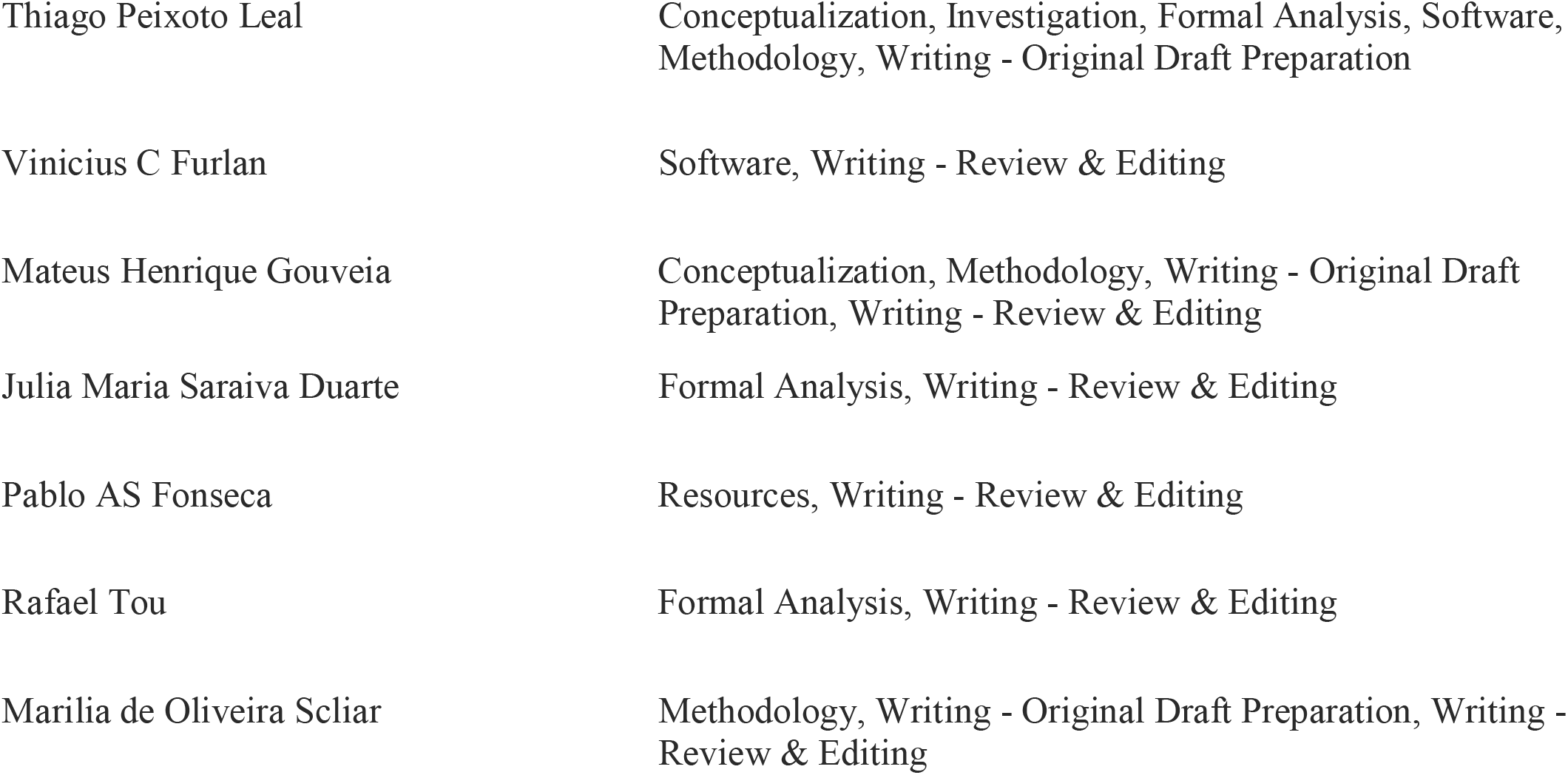
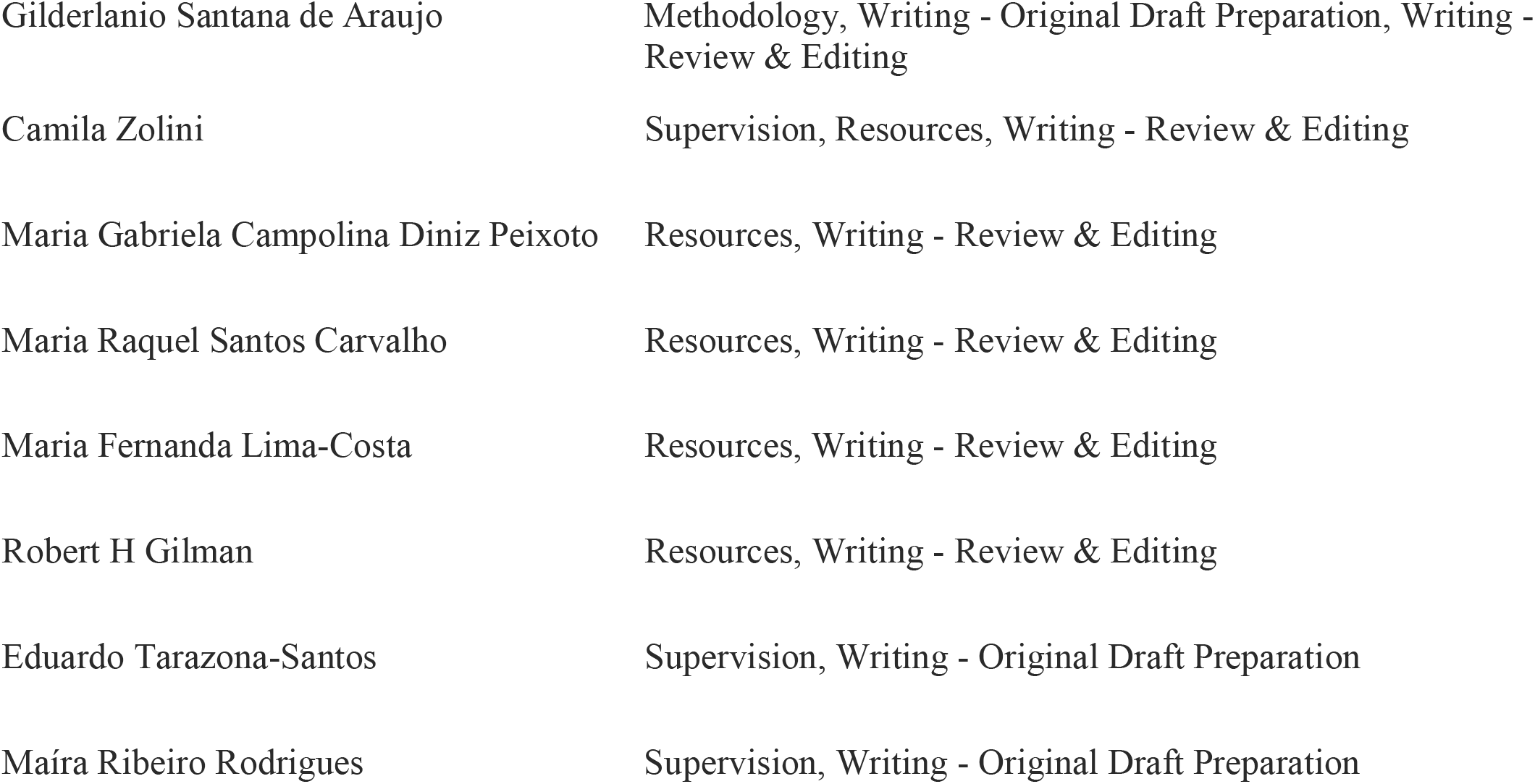

## Acknowledgements

Not applicable

