## Supplementary material for "NAToRA, a relatedness-pruning method to minimize the loss of dataset size in genetic and omics analyses": SI

### Paper: NAToRA, a relatedness-pruning method for minimal reduction of dataset size in genetic and omics analyses

**Sections**

#### S1 Relatedness and Population genetics

In this work we use the population genetics concept of identity by descent (IBD) to define genetic relatedness. Two individuals are considered genetically related if they have identical alleles by descent, i.e., these alleles are derived from a recent common ancestor. Specifically, the kinship coefficient ($\Phi_{i,j}$) represents the probability of two alleles, randomly sampled at the same locus, of two individuals i and j being identical by descent [1,2].

The kinship coefficient can be expressed as a function of the probabilities of identity by descent (IBD), denoted by equation $\Phi_{i,j} = \frac{\delta^1{}_{ij}}{4} + \frac{\delta\ {}^2{}_{ij}}{2}$, where $\delta^1{}_{ij}$ is the probability of individuals i and j sharing one IBD allele and $\delta\ {}^2{}_{ij}$ is the probability of i and j sharing two IBD alleles [3,4]. The PI_HAT, another metric used in this work, is calculated by equation $PI\_HAT = \frac{\delta^1{}_{ij}}{2} + \delta\ {}^2{}_{ij}$. Table S1 shows the kinship coefficients corresponding to specific pairs of relatives.

There are different methodologies for estimating the kinship coefficient between two individuals through genetic data when the genealogical relationships are not known. In this work we use two methodologies: (i) KING [5], that calculates the kinship coefficient and (ii) PLINK [6], that calculates the PI_HAT. In addition, we can calculate the kinship coefficient using REAP (Relatedness Estimation in Admixed Populations)[3,4]. PLINK is more suitable to measure relatedness in populations that are not admixed and REAP is more appropriate for admixed populations.

Also, since kinship values inferred by the above methods do not always match the theoretical values, some tools use the geometric mean of theoretical values to define the interval that represents the desired relationship degree [5], that is, given the theoretical values of first (0.25) and second degree (0.125), they use the geometric mean ($\sqrt{0.125\ x\ 0.25}$) between these two numbers to define the value that separates the non-theoretical values between first and second degree.

For population genetics analyses that require independent observations, as opposed to allowing related individuals as part of the dataset, a common approach is to prune out all related individuals or remove related individuals randomly [7];[8,9]; [10]. This is to avoid confounding factors in such analyses. For instance, in [3], while performing ADMIXTURE analysis, the authors found a biogeographical cluster exclusive to the Bambui population (K=7). Further investigation showed that these clusters were related individuals that the algorithm incorrectly inferred to be a biogeographical cluster (Figure S1). Although the above pruning strategies can be used for a wider range of analyses, they can lead to unnecessary dataset loss. In this work, we call “general strategy” this first strategy of pruning all individuals that have at least one relationship greater than a cutoff value of Kinship < α (α is the cutoff value).

#### S2 Implementation of NAToRA

NAToRA (Network Algorithm To Relationship Analysis) is a methodology that models the genetic relatedness of datasets as a complex network to resolve the problem of reducing relatedness in a population dataset while minimizing the number of excluded individuals. NAToRA was implemented as a greedy algorithm and is based on Complex Network Theory. It was implemented in Python using the library NetworkX [11].

The input data is a pairwise relatedness matrix or Genetic Relationship Matrix (GRM) and a relatedness cutoff value that indicates the maximum kinship degree that should be accepted between the individuals in the matrix. The relatedness metric used in this matrix can be of any kind. In this work we use the kinship coefficient ($\Phi_{i,j}$)and the The PI_HAT, however, the user may input other metrics. Fonseca et al. in [12] used G-Matrix as input for NAToRA.

From the GRM, the algorithm creates a network $N$ (Figure S2 (a)). In the context of Graph and Complex Network Theory, a network G (or graph) is a pair G=(V,E) consisting of a set of vertices V connected by a set of edges E [13]. In our representation, the vertices are individuals and there is an edge between two individuals if the genetic relationship is nonzero. Since kinship is a bi-implication relationship (if *a* is relative of *b* then *b* is relative of *a* with the same degree), our network model is undirected and weighted by the values of GRM. In this work, all concepts of Complex Network and Graph Theory are for undirected networks [13]; [14].

In our methodology we allow the user to select the minimum relatedness degree to be present in the dataset (the relationship cutoff α). The network that only has edges with value above the cutoff value is called $N_c$(Network with cuts) (Figure S2 (b)). Using network $N_c$ the algorithm performs two analyses: (*i*) Families detection and (*ii*) iterative pruning of individuals based on node centrality metrics.

To detect families, the algorithm identifies all connected components of Network $N_c$. A connected component is a set of nodes that has at least one path (sequence of edges which connect a sequence of nodes) between all of them. A connected network is a network that has a path between all nodes and it has only one connected component (Figure S3(a) is a connected network). On the other hand, a network is disconnected when there is at least one pair of vertices that are not a path by any path. This analysis is shown in Figure S2 (b) through the colors of the nodes, in which each color represents a different family.

After identifying the families, the algorithm creates a file with this information because it can be used, for example, as PLINK's family ID (FID) or as a categorical co-variable in association studies. After creating this output, the algorithm starts the iterative pruning of individuals based on the node centrality metrics. This step tries to exclude the minimum number of individuals from the dataset. The problem to exclude the minimum number of individuals to obtain a network without edges is analogous to finding the maximum clique in the complement network, which is an NP-Complete Problem.

In the context of graph theory [13], a complement network H of a network G is a pair H=(V,B), where B is a set of edges that connect two nodes *u* and *v* if and only if there is not an edge connecting *u* and *v* in network G. In this modelling, the complement network is the network of non-related individuals(Figure S3). A clique of a network G=(V,A) is a subnetwork which for each pair of nodes (*u,v*) there is an edge between them (Figure S4). Finding the maximum clique, *i.e.*, the clique that has the largest number of nodes, is an NP-Complete Problem [14]. In our model, to find the maximum clique in the complement network is to find the largest set of individuals which all are mutually unrelated. Because it is an NP-Complete problem, which can be computationally unfeasible depending on the network, we implement a heuristic that consists of the iterative pruning of individuals based on centrality metrics.

In complex networks theory, centrality metrics aim to associate a value of importance to a node and/or edge, that is, the more central a node and/or edge the more important it is to the network. Each centrality metric has its own criteria to defining the importance of a node and/or edge [13].

In this work we tested three node centrality metrics: (i) node degree centrality, calculated by the number of edges connected to the node;(ii) node betweenness centrality, based on the shortest path, and it is a measure of the number of shortest paths that pass through this node (the more paths pass through a node, the greater its centrality; and (iii) closeness centrality, which is the mean distance from a node to the other nodes [13].

The node betweenness centrality and the node closeness centrality assign higher values of centrality to nodes that have edges with small values, but in our methodology we want to reduce the relatedness in the dataset, represented by higher values of edges. For this purpose, we used a parameter β in which the user tells to the algorithm the highest possible value of the pairwise relatedness matrix (for example, using the kinship coefficient we set β as 0.51) . With this value we calculate a new value by subtracting β with all values present in the matrix. In this way, the individuals with the highest degree of relationship will have the smallest edges in the network, which will make them the most central nodes in the network.

For each family identified by the algorithm, it performs the following steps: (*i*) calculates the centrality metric for each individual and (*ii*) prunes the most central individual, storing the ID. If there is a tie in the most central individual, we prune the individual that has the higher sum of the weights of the remaining edges (or the smallest if the edges were subtracted by β). Steps (*i*) and (*ii*) are repeated until only individuals without edges or pairs of individuals linked by only one edge are kept in the dataset (Figure S2 (c-h)).

When there are only pairs of individuals connected in the network, the elimination based on centrality metrics loses efficiency since both nodes will have the same centrality value. To solve this, we implemented a tiebreaker that consists of calculating the centrality of each individual in the pair using network N using selecting all edges between an interval (min and max tiebreaker value, which default value is 0.0221 an the relationship cutoff α) instead of network Nc, and excluding the most central individual (Figure S2 (i)(j)).

In Figure S2 we present an example of how the NAToRA works and all steps of NAToRA can be seen in Figure S5. Because the node betweenness centrality and node closeness centrality did not work properly (see Results section), we do not represent normalization in the flowchart.

#### S3 Genealogy simulator

To generate simulated data we implemented a Genealogy Simulator. This simulator aims to create genealogy with reproductive behavior similar to expected in human populations based on parameters provided by the user, allowing to create several different scenarios. In our model, we defined that full siblings can not have children and it is only possible to have offspring between individuals of the same generation. After generating the genealogy, the algorithm calculates the theoretical kinship coefficient (Table S1) among all pairs of related individuals.

The simulator was implemented following the divide and conquer paradigm, which consists of dividing a given problem into two or more smaller problems that will be solved independently using the algorithm defined and later combining these solutions in order to produce a solution to the original problem [14]. The problem of the simulator is to generate a genealogy of a total G generations, so that each generation has 4 parameters defined by the user with the model restrictions explained earlier. In order to solve this, we divided into sub-genealogy (generation 1 with 2, generation 2 with 3, …, generation G-1 with generation G) and, after generating all sub-genealogies, we joined each of them to create a genealogy with all generations, correcting eventual inconsistencies, ie, siblings in generation *k* being parents of an individual in generation *k+1* (Figure S6).

The algorithm works with 4 parameters: Number of male and female of generation g ($N_{w,g}$ and$N_{m,g}$, respectively), proportion of unrelated in generation g ($P_{u,g}$) and proportion of half-brothers in generation g ($P_{s,g}$) (Figure S6(a)). Despite all the generations having the 4 parameters, we consider the first generation (founders) are all unrelated between them, ie, they do not have a common ancestor in the genealogy generated.

With all parameters, the algorithm creates G-1 sub-genealogies of continuous generations, denoted by $F_{k,k+1}$, where k is a generation different from G. In other words, we created a sub-genealogy with generations 1 and 2 ($F_{1,2}$); a sub-genealogy with generations 2 and 3 ($F_{2,3}$); … ; a sub-genealogy with generations G-1 and G ($F_{G-1,G}$).

To generate the sub-genealogy to a generation k with generation k+1 ($F_{k,k+1}$), the first step is to select which individuals of generation k+1 that will not have kinship in this sub-genealogy, that we call $U_{k+1}$ (Unrelated in generation k+1). These individuals are not the progeny of any individual of k and, to select this individuals, we use the proportion of unrelated in generation k+1 ($P_{u,k+1}$), calculating the number of individuals through equation $A_{u,k+1} = P_{u,k+1} * (N_{w,k+1} + N_{m,k+1})$. Because the number of individuals is an integer number, we round $A_{u,k+1}$ and select randomly the individuals ( $U_{k+1}$) to not have kinship in subgenealogy $F_{k,k+1}$ (Figure S6(b) and (i)).

After the selection of individuals belonging to $U_{k+1}$, we create the maximum of couples of individuals randomly in generation k (Figure S6 (c) and (j)) and distribute, randomly, the individuals who were not selected to be the progeny of the pairs (Figure S6 (d),(e),(f),(k)(l) and (m)).

After assign progeny to the couples, the algorithm calculates the minimal number of half-siblings $A_{s,k+1}$, calculated by the equation $A_{s,k+1} = P_{s,k+1} * (N_{w,k+1} + N_{m,k+1})$. Because the number of individuals is an integer number, we round $A_{s,k+1}$ (Figure S6 (g) and (n)) and, if the number is greater than 0, we replace one of the parents randomly to generate half-siblings (Figure S6 (h)) . This process is repeated until we have at least $A_{s,k+1}$ half-siblings (Individual 8 in Figure S6 (h)).

In short, we create all possible parents in k and assign children to them (and some pair may not have children), keeping unrelated the $U_{k+1}$ selected individuals. If the sub-genealogy $F_{k,k+1}$have half-siblings, the algorithm replaces one of the parents until have at least $A_{s,k+1}$ individuals. After these steps we have the sub-genealogy $F_{k,k+1}$ (Figure S6 (h)(m)).

After the generation of all sub-genealogy ($F_{1,2}$, $F_{2,3}$… $F_{G-1,G}$), the algorithm merge them based in chronological order, ie, first merge $F_{1,2}$ with $F_{2,3}$, after merge with $F_{3,4}$, .., and finally with $F_{G-1,G}$. To achieve this goal, the algorithm maps the progeny $F_{k,k+1}$ as the parents in $F_{k+1,k+2}$(Figure S6 (o)). After each merge, the algorithm looks for possible inconsistencies, for example, two individuals are a couple in $F_{k+1,k+2}$ and be siblings in $F_{k,k+1}$. In this case, the algorithm changes this individual randomly until there is no more inconsistency.

***Simulated scenarios***

Using the simulator described we generate four scenarios with four generations each:

- Scenario 1: 10 men and 10 women for each generation (total of 80 individuals)
- Scenario 2: 50 men and 50 women for each generation (total of 400 individuals)
- Scenario 3: 100 men and 100 women for each generation (total of 800 individuals)
- Scenario 3: 500 men and 500 women for each generation (total of 4000 individuals)

In all scenarios the proportion of half-siblings is 0 and the proportion of unrelated is 0.5 for all generations. We generate 100 genealogies (total of 400 simulated tests). Since most of the steps are randomized, we expect each execution to generate different scenarios. With this data we evaluate the performance of the algorithm for genealogies that differ in topology and in the number of individuals that compose them. With this data, we evaluate the performance of the algorithm for genealogies with different numbers of individuals. Since genealogies are generated in a non-deterministic way, we repeat each "case" several times and performance evaluation considers an average of these cases (Figure S10).

#### S4 NAToRA Tests with Simulated Scenarios

We used the genealogy simulator to simulate four scenarios with different datasets sizes. For each scenario, we simulate 100 genealogies that were used to compare the centrality metrics.

To obtain the optimal result we select all connected components and generate the complementary network for each connected component. We use the "find_cliques" function implemented in NetworkX library, that returns a list with all cliques with a computational cost O($3^{\frac{n}{3}}$). The result from clique is a list of unrelated individuals to keep. This output is different from the heuristics based on centrality metrics, which is a list of individuals to be removed.

The comparison was made by the difference between the number of individuals to be eliminated using a centrality metric (heuristic) and the number of individuals to be eliminated by the optimal result. The best result will be 0, which means that the number of individuals to be eliminated by heuristic is the same as the number of individuals eliminated by the optimal algorithm (clique). All genealogies generated by the simulator have different configurations and therefore the individuals to be pruned in each genealogy will be different, making impossible to compare the different simulations for each scenario.

Using Scenario 1 the heuristics based in node degree centrality and node betweenness centrality obtained very similar results (Figure S10 (A)). However, we noticed that the node degree centrality has the best results when analysing the other three scenarios (Figure S10 (B), (C) and (D)), presenting many results with the same number of eliminated individuals as optimal algorithm for all scenarios. The heuristic based on node closeness centrality was less efficient in all tests, showing that this metric is not ideal for reducing relatedness in a dataset.

In addition to obtaining better results, node degree centrality presented shorter execution times when we compared the execution times present in Table S2, Table S3, Table S4 and Table S5 (which considers only processing time, that is, does not consider such as scheduling, I / O requests, etc) of the tests that have been completed. Some tests (2 in scenario 3 and 13 in scenario 4) did not complete after 7 months running, so they were not taken into consideration.

#### S5 Genetic Datasets

To test out algorithm with real data, we used three databases: (i) The Bambuí Aging Cohort Study (BAMBUÍ, admixed individuals), (ii) Matsiguenkas indigenous of Amazon Yunga, the rainforest transitional region between the Andes and the Lower Amazonia, (SHIMAA) and (iii) Cattle from National Breeding Program for milk production of Guzerá cattle breed (GUZERÁ). We chose a mix of human and non-human datasets to show the general applicability of the method and because non-human datasets usually have higher inbreeding and, thus, are more affected by relatedness-pruning.

The Bambuí dataset is composed by admixed individuals of Bambuí Aging Cohort Study situated in Minas Gerais, Southeast Brazil [3]. This dataset integrates the EPIGEN-Brazil Project, in which 1,442 individuals were genotyped for 2.2 million SNPs (Illumina 2.5M). We calculated the kinship coefficient using REAP software [4] (Figure S7) and, unlike the other EPIGEN-Brazil cohorts, this dataset includes several families with 5 individuals or more with first or second kinship degree($\Phi_{i,j} > 0.10$).

The Shimaa dataset is composed of 43 individuals [15]. To analyse the relationship structure of this dataset, we calculated the kinship coefficients using PLINK software. We observed 17 families when using second-degree cutoff ( $\Phi_{i,j} > 0.0884$) and individuals have an average of 4.22 relationships above second degree (sd = 4.76) (Figure S8).

The Guzera dataset is composed of 1,036 Guzerá (Bos indicus) cattle from the National Program for Improvement of Guzerá for Milk, genotyped using Illumina Bovine SNP50 v2 Bead Chip. This dataset was selected due to its very particular genetic composition and it is part of a Multiple Ovulation Embryo Transfer (MOET) breeding program. Typically, MOET programs are based on the selection of top individuals from both sexes to include in the crosses. Superovulation is induced in donor cows, embryos were produced in vitro, and transferred into receptor cows. In addition, the number of bulls included in the breeding program is even smaller than the number of cows. As a consequence, MOET individuals are composed of a large number of both maternal and paternal half-sibs, producing extremely complex, hard to disentangle pedigrees. We calculate the kinship coefficient using PLINK software (Figure S9) and we find that the majority of individuals are in a single family ($\Phi_{i,j} > 0.0884$) and the others are present in other minor components and we observed only 15 non-related individuals. In this dataset there are six families and the individuals have an average of 34.53 relationships above the second degree (sd = 47.33)

#### S6 NAToRA Tests with Genetic Datasets

As observed in simulated data, the heuristic based in node degree centrality obtained the best results and the shortest execution time (Table S6). In some cases, the optimal algorithm obtained results in computational time close to the node degree centrality. This low computational time was achieved because the network has a high edge density and low number of individuals per family, which results in a complementary network with few edges and few nodes, thus reducing the search space for the maximum clique. The optimal result for GUZERÁ was not concluded after seven months of execution and, therefore, was not considered in the analysis.

#### S7 Comparison with existing methods

In some studies, in which it was necessary to control the relationship between individuals in the dataset, researchers implement an algorithm that we will call here general strategy or Kinship < α. In this technique the researchers infer the relationship between the individuals and remove all individuals that have at least one relationship greater than a cutoff α, keeping only individuals with relationships less than α value ([10]. We used this strategy to show the worst case of relatedness reduction.

The software PLINK [6], one of the main bioinformatics tools used in genomic analysis, also provides a relatedness-pruning filter through the parameter "--rel-cutoff". For this analysis, the software calculates the PI_HAT (values range from 0 to 1), selects the relationship that has a value greater than a cutoff chosen by the user and excludes one of the individuals. After the processing the software makes available the list of individuals to be removed.

Another software that implements relatedness-pruning is KING [5] using the parameter "--unrelated --degree <degree chosen by the user>". For this exclusion the software calculates the kinship coefficient for all pairs of individuals and clusters those with kinship greater than the value selected by the user (they consider this cluster as family). After that, the individuals of the family are ranked based on the count of unrelated family members (i.e. the number of individuals that kinship coefficients estimated are lower than the value selected by the user). The algorithm creates the unrelated dataset selecting the individuals with highest rank and, if this individual is not related to any previous individuals, this individual is included in the unrelated dataset.

To compare our methodology with PLINK, KING and the general relatedness-pruning strategy we use the three real databases described above (BAMBUÍ, SHIMAA and GUZERÁ). Although NAToRA accepts a variety of relatedness metrics as input, we cannot directly compare PLINK's and KING's relatedness-pruning methods due to their different input metrics (PI_HAT for PLINK and kinship coefficient for KING). Due to the fact that there is no way to perform the relationship pruning without having to make the inference, the relationship network for PLINK and KING will be different, making it not directly comparable. Thus, we performed groupwise comparisons between each one of these methods, NAToRA, and the general relatedness-pruning strategy, using the equivalent metrics for excluding individuals with second-degree kinship or higher. We calculate the relationship matrix through PLINK and KING for all databases and use these values to perform the experiments. In the KING experiment we used the second degree cutoff (0.0884 according the manual, which is the geometric mean between the second and third degree kinship theoretical value ) and in PLINK comparison the cutoff value chosen was 2x the KING cutoff value (0.1768) because the PI_HAT is 2x the kinship coefficient (Cormack, Hartl, and Clark 1990). In the experiment using the real dataset, we measured the number of individuals and of relationships above the cutoff that remained in the dataset after the relatedness pruning. The desirable result will have the highest number of individuals and the lowest number of remaining relationships above the cutoff. The results of the comparison between PLINK, KING, NAToRA and the general strategy are presented in Table S7. We used the Brazil EPIGEN dataset (that is composed of the BAMBUI dataset and two other cohorts from EPIGEN-Brazil project [3]) to evaluate the performance of NAToRA with datasets with more individuals.

When we compare the remaining individuals after the pruning process, NAToRA excludes more individuals when the Network Family has many individuals and a high edge density. We expect this behavior since NAToRA aims to exclude all relationships greater than the specified cutoff and the other methods are not so strict on this point. If the analysis demands that there is minimal individual loss, we recommend increasing the cutoff values (the number of individuals to be removed can be estimated by the flag “--test <highest possible value of metric used>”, Figure S17).

In this work we presented and compared the performance of NAToRA with two popular methods, the relatedness-pruning filters implemented in softwares PLINK and KING, and a general relatedness-pruning strategy found in literature. We show that NAToRA outperforms all methods by removing all relationships above the desired kinship cutoff from three real datasets and preserving more individuals in most tested scenarios.

For the Guzerá breed dataset, particularly, the number of individuals excluded using all tested software was large. This was expected since GUZERÁ is a dataset with known high relatedness. In a context of high relatedness, NAToRA was able to better preserve the alleles of lower frequency, generating a dataset that better represents the allele composition of the original dataset.

#### S8 The impact relatedness-pruning in genetic diversity and MAF distribution

After obtaining datasets pruned by relatedness above a selected cutoff through NAToRA, PLINK, KING and the general relatedness-pruning strategy, we evaluated the impact of these methods on some characteristics of the final datasets. Specifically, on the distribution MAF (Minor Frequency Allele) classes and on dataset diversity through PCA (Principal Component Analysis). This verification is important because our approach is based on excluding individuals, which can generate sub-datasets with different characteristics from the original dataset in addition to maximizing the reduction of inbreeding and minimizing the loss of dataset size.

For PCA we used the software EIGENSTRAT that uses genotype data to infer continuous axes of genetic variation. The main objective of the algorithm is to reduce the variation to a small number of dimensions describing as much variability as possible [16].

The PCA was executed with four databases for SHIMAA, BAMBUÍ and GUZERÁ: (i) original database, (ii) without individuals which has relationship greater than 2nd degree (Kinship < 2nd degree), (iii) without individuals removed by NAToRA and (iv) without individuals removed by tool that infer the relationship (PLINK or KING). Using PCA analysis with all subsets of data we can assess the impact of using NAToRA on the axis capturing most of genotyping variance (PC1 and PC2) on the global data variability analysing regions in the diversity planes that will be unsampled due to individual exclusion.

In addition to PCA analysis, we conducted a NAToRA impact study on MAF frequencies, which represent the frequency of the second most common allele. The MAF distributions are shown in Figure S11. In this analysis we observed that removing all related individuals (Kinship < 2nd degree) has a major impact on MAF frequencies, especially in the monomorphic category (data that was not monomorphic being monomorphic in the pruned dataset). We did not observe a large difference between the MAF distributions created by the heuristic based on NAToRA and PLINK or KING.

Analysing the PCA results, we observed that BAMBUI, SHIMAA and GUZERÁ PC1 and PC2 (Figure S12, Figure S13 and Figure S14) have some regions without individuals in the database Kinship < 2nd degree, showing that this strategy has a big loss of variability. This problem is more compelling in the GUZERÁ dataset with relationships inferred by PLINK, in which individuals are present in a small portion of the plan. This is an expected issue because the Kinship < 2nd degree has fewer individuals in all datasets. The NAToRA methodology maintains a large part of the variability in all analyzes, showing a better or comparable performance to PLINK and KING, which do not guarantee the removal of the entire relationship from the dataset. To facilitate the visualization of this issue (variability vs number of individuals vs number of remaining relationships) see the Figure S15.

In addition to the results of this paper, Fonseca et al. [12] showed that NAToRA contributes to reducing GWAS inflation values using simulated and cattle herd data in the same GUZERÁ dataset. With these analyses we can say that NAToRA, either the heuristic version or the optimal algorithm, reduces kinship without much loss of genetic diversity.

#### S9 The NAToRA pipeline used in Sample Database - a case study

To exemplify the use of NAToRA and all supporting scripts we use the Sample Database (n=1,442) with three tools: KING, REAP and PLINK. The calculation and the conversion to NAToRA input format (adjacency list) is unique for each tool, but after this conversion, all the other steps are the same. You can find the conversion scripts used below on the NAToRA website (http://ldgh.com.br/natora/) under the User Guide tab or https://github.com/ldgh/NAToRA_Public.

##### To calculate kinship coefficients using KING

**The first step** is to calculate the kinship coefficient using KING, with the command line shown in (1).

*king --kinship -b SampleFile.bed --prefix Sample_KING* (1)

When calculations are done, the **second step** is to convert to NAToRA input format (using the command line (2)), that is an adjacency list. We used the .kin0 file because this file contains all kinship coefficients calculated by KING.

*perl KING2NAToRA.pl --input SAMPLE_KING.kin0 --output SAMPLE_KING* (2)

##### To calculate PLINK's PI_HAT metric

PLINK calculates the PI_HAT metric through the flag --genome, (note that PI_HAT is twice the coefficient kinship presented in this paper). This metric calculation requires a LD-pruned dataset. Thus, the steps to generate the PI_HAT matrix are:

*plink --bfile SampleFile --indep-pairwise 200 25 0.4 --out LD* *(3)*
*plink --bfile SampleFile --extract LD.prune.in --out SampleFile_LD --make bed* *(4)*
*plink --bfile SampleFile_LD --genome --out Sample_PLINK* *(5)*

When calculations are done, the **second step** is to convert to NAToRA input format (command line (6)), that is an adjacency list. In order to allow all steps of the pipeline, we convert PI_HAT to kinship coefficient with --kinship flag in PLINK2NAToRA.pl

perl PLINK2NAToRA.pl --input Sample_PLINK.genome --out SAMPLE_PLINK --kinship *(6)*

The flag --kinship is used to indicate to the algorithm that it must calculate and output the kinship coefficient using IBD1 and IBD2.

##### Common steps (Online Webtool)

After we convert the output file from kinship estimator tools, **the next step** is run NAToRA uploading the generated file at http://ldgh.com.br/natora/. The file can be a plain text file or a zipped file (.zip or .rar) if you have larger files or connection issues (Figure S16). The algorithm parameters can be selected by clicking on the gear button (Figure S17), as follows:

**Use kinship** (optional): option that can be used if your relationship metric is the Kinship coefficient. This option will automatically set the mandatory parameters with default values based on the kinship degree selected.
**Generate sets** (optional): this option will generate multiple sets of unrelated individuals and automatically prompt a download window for the generated files (see Section S10).
**Elimination method** (optional): possible elimination methods are the Natora option, which will check your network and choose automatically between optimal and heuristic elimination; optimal or heuristic. ATTENTION: You should be careful in selecting the optimal version for large datasets, since it can be computationally intractable (see section S11).

**Min tiebreak value** (mandatory): The minimal relatedness value used to generate the network that will be used in the tiebreak process (see section S2).

**Max tiebreak value** (mandatory): The maximal relatedness value used to generate the network that will be used in the tiebreak process (see section S2).

**Cutoff** (mandatory): A relatedness cutoff value indicating the maximum relatedness degree expected in the dataset after the pruning process.

If your relatedness metric is different from Kinship Coefficient, such as PI_HAT, you should manually set the mandatory parameters above. It is important to note that these mandatory values must be of the same type of the relatedness metric in your input file (e.g., PI_HAT values range from 0 to 1; kinship coefficient values range from 0 to 0.5). After that, you can click the Run button and the analysis window will show after the upload.

After the upload process, the webtool will show the result screen. In this screen the window is splitted in three sections: (i) the input (left), (ii) the network view (center) and (iii) the output (right) (Figure S18).

In the left section you can see the parameters used in the analysis and change it if you want to redo the analysis without needing to upload the data again. To do this, you can change the values and click at "Generate Network". You can also download the GML file ("Download .GML file"), a file that is the input file to software that analyse and plot network data. When we click on this button, the webtool will download a file named GraphData.gml. We used yEd Graph Editor (https://www.yworks.com/products/yed) to plot the network (Figure S19). In this section there is also the Input Network Information with the number of individuals and relationships present in the network.

The section presented in the middle is the network view section. It allows the user to visualize the input and output data as a Network.

The right section is the output section, where the user can download the Family List, a file with the individuals and which network family they belong to, and the Removed individuals list, a file with the list of individuals that should be removed from the dataset. It also shows the output network information, with the number of individuals and the number of remaining relationships.

If the user is using PLINK to analyse data, we provide a script (command (7)) that converts NAToRA's output file *removedNodes.txt* into a compatible format (i.e., <Family ID> <Individual ID>) to use with PLINK's --remove flag (command (8)).

*perl generatePLINKlistToExclude.pl --fam SampleFile.fam --remove removedNodes.txt --out listRemove* *(7)*

*plink --bfile SampleFile --remove listRemove --out SampleFile_withourelatives --make-bed* *(8)*

After this step, we have the database without any kinship coefficient larger than 0.0884.

#### Common steps (Local running)

After we convert the output file from kinship estimator tools, **the next step** we run NAToRA with flags "--test 0.5 --kinship" to infer the number of sample loss by cutoff (command line (7)). The "--test 0.5" indicates to NAToRA to calculate the sample loss from 0.05 until 0.5 and "--kinship" will use the kinship coefficient as reference (Figure S20(A), kinship intervals are shown in Figure S20(B)). The algorithm returns to the user the min, theoretical and max values for each relationship degree (Figure S20(C)).

*python NAToRA_Public.py --input BAMBUI_NAToRA --test --kinship* *(7)*

Using the data in NAToRA format (<ind1> <ind2> <value>), the **next step** is to use the script generateGML.pl to plot the families network in .gml format (command line (8)). In our example we consider as relatives individuals with kinship coefficient bigger than 0.0884. This value is used to KING to specify a second degree or greater.

*perl generateGML.pl --input BAMBUI_NAToRA --cutoff 0.0884 -d --out BAMBUI.gml* *(8)*

GML (graph markup language) is a format of file accepted by software that work with complex networks. To plot this result we used yEd Graph Editor (https://www.yworks.com/products/yed) (Figure S21).

To remove all kinship larger than second degree we can use the min value returned by algorithm, ie, 0.0884 (9) or --degree 2 (10). After running, the algorithm will generate the file<prefix>_toRemove.txt, in which the <prefix> is defined by the "-o" parameter. The parameters -v and -V are optional and we will use them to demonstrate its use (parameters used in the refined exclusion, as explained in Implementation of NAtoRA section).

*python3 NAToRA_Public.py --input BAMBUI_NAToRA -c 0.0884 -o NAToRA_BAMBUI -v 0.03 -V 0.2* *(9)*
*or*
*python3 NAToRA_Public.py --input BAMBUI_NAToRA --kinship --degree 2 -o NAToRA_BAMBUI -v 0.03 -V 0.2* *(10)*

Using the file *NAToRA_BAMBUI_*toRemove.txt you can exclude the individuals using PLINK. To do this, we use the script *generatePLINKlistToExclude.pl* (command (11)) to create the list with the format used in PLINK(<Family ID> <Individual ID>) to use the parameter --remove (command (12))

*perl generatePLINKlistToExclude.pl --fam BAMBUI_Autosomic_SmartCleaning.fam --remove NAToRA_BAMBUI_*toRemove.txt *--out listRemove* *(11)*

*plink --bfile BAMBUI_Autosomic_SmartCleaning --remove listRemove --out BAMBUI_withourelatives --make-bed* *(12)*

After this step, we have the database without any kinship coefficient larger than 0.0884.

#### S10 Using NAToRA to create multiple sets of unrelated individuals

As seen in NAToRA Tests section, our method is able to remove relationships keeping the number of individuals removed very close to the minimum possible (optimal algorithm). Nevertheless, there are some analyses in which it is necessary to remove kinship without losing individuals, to reduce the statistical power of the analysis. For analysis that we can divide the data into subsets without affecting the final result, we can use NAToRA several times to get a total of R sets of unrelated individuals, in order to perform independent analysis with each of the R sets of unrelated individuals.

The first step to create the R sets of unrelated individuals is to select in the $N_c$ network all unrelated individuals and store this list of individuals. In this work we will call this list DU (Dataset Unrelated) and we use DU as a quality control set. After that, we run NAToRA normally and the algorithm will return a list of individuals to be removed. After we eliminate individuals we obtain the first set of unrelated individuals, that we will call $AD_1$ (Analysis Dataset 1).

After we create $AD_1$, the algorithm removes all individuals present in $AD_1$from the input database for NAToRA. After removing these individuals, the algorithm reinserts the DU (creating a new input data) and runs NAToRA again, generating a new list of individuals to be deleted to obtain an unrelated dataset above the cutoff value. We will create $AD_2$removing the output list from NAToRA from the input database for the last run of NAToRA (not the original database).

The algorithm to create multiple datasets of unrelated individuals consists in to remove from original dataset individuals present in any previous analysis ($AD_1$, $AD_2$, …), to add the DU individuals, to execute NAToRA and to remove from the last input dataset all individuals present in the list output by NAToRA. The algorithm is finished when the output from NAToRA is empty,i.e., the input to NAToRA has no relationship. All steps are presented in Figure S5(D) and there is an example in Figure S21.

As each set is independent, this strategy allows to perform analyses, such ADMIXTURE [17], for all dataset without the problem of sample loss and avoiding the problems that kinship can cause. The user can compare the different runs through individuals present in DU. This approach was used in [18].

You can perform this analysis using the “Generate Sets” option in the gear button. When selected the webtool will download, automatically, a list with individuals and the set that they should be present. With this information the user can create multiple genetic files with which all individuals are present without any relationship.

#### S11 Identifying feasible features for running the optimal algorithm

After comparing the heuristic and optimal algorithm, we found out that for some cases it was feasible to use the optimal algorithm (Table S2, Table S3, Table S4, Table S5, Table S6). In this section we conducted studies with simulated data to try to identify which network characteristics allowed the use of optimal algorithm.

In these tests we used a generator of small world network implemented in NetworkX library to simulated networks with different number of nodes (100, 200, 300, 400, 500, 600, 700, 800, 900, 1000), with different amounts of edges per node (10%, 30%, 50%, 70% , 90% of total possible edges calculated by the formula $Total = \frac{n*(n-1)}{2}$) and with the probability of 50% of rewiring each edge to create different networks with the same parameters. Using this approach, we can generate families with different amounts of individuals and family relationships. For each combination of number of nodes and the amount edge per node, we generate 10 networks and execute the optimal algorithm and the time distribution is shown in Figure S22.

After these analyses we conclude that the scenarios, in which it is feasible to use the optimal algorithm are networks whose largest family have few individuals (<100 individuals) or with a high edge density (> 50% of the total possible). These conditions make sense considering how we perform the analysis with the optimal algorithm, in which we generate the complementary network and use the clique detection algorithm. A complementary network of a network with a high edge density has few edges having a small search space.

With this information we implemented a feature that the algorithm will choose between heuristic and optimal mode based on network features previously explained (NAToRA in the elimination method section). The user can choose the elimination method manually by selecting the heuristic version or the optimal version in the elimination method section.

11. Hagberg AA, Schult DA, Swart PJ. Exploring Network Structure, Dynamics, and Function using NetworkX. In: Varoquaux G, Vaught T, Millman J, editors. *Proceedings of the 7th Python in Science Conference*. Pasadena, CA USA; p. 11–5.

12. Fonseca PAS, Leal TP, Santos FC, Gouveia MH, Id-Lahoucine S, Rosse IC, et al.. Reducing cryptic relatedness in genomic data sets via a central node exclusion algorithm. Molecular Ecology Resources.

13. Newman M. Networks: An Introduction. Oxford: Oxford University Press; 2010.

14. Cormen TH, Leiserson CE, Stein RLREC. Algoritmos: teoria e prática. Elsevier Editora; 2012.

15. Borda V, Alvim I, Aquino MM, Silva C, Soares-Souza GB, Leal TP, et al.. The genetic

structure and adaptation of Andean highlanders and Amazonian dwellers is influenced by the interplay between geography and culture. *PNAS*. 2020; doi: 10.1073/pnas.2013773117.

16. Price AL, Patterson NJ, Plenge RM, Weinblatt ME, Shadick NA, Reich D. Principal components analysis corrects for stratification in genome-wide association studies. *Nat Genet*. 2006; doi: 10.1038/ng1847.

17. Alexander DH, Novembre J, Lange K. Fast model-based estimation of ancestry in unrelated individuals. *Genome Res*. 2009; doi: 10.1101/gr.094052.109.

18. Lima-Costa MF, Mambrini JV de M, Leite MLC, Peixoto SV, Firmo JOA, Loyola Filho AI de, et al.. Socioeconomic Position, But Not African Genomic Ancestry, Is Associated With Blood Pressure in the Bambui-Epigen (Brazil) Cohort Study of Aging. *Hypertension*. 2016; doi: 10.1161/HYPERTENSIONAHA.115.06609.
