## Supplementary material for "NAToRA, a relatedness-pruning method to minimize the loss of dataset size in genetic and omics analyses": SI Figures and Tables

**Table S1.** Theoretical values $\delta_{i,j}^{0}$ (Probability of IBD = 0), $\delta_{i,j}^{1}$ (Probability of IBD = 1), $\delta_{i,j}^{2}$ (Probability of IBD = 2) and $\Phi_{i,j}^{\blacksquare}$ (Kinship coefficient) for each kinship degree (Thornton et al. 2012)

| Outbred relationship | $\delta_{i,j}^{0}$ | $\delta_{i,j}^{1}$ | $\delta_{i,j}^{2}$ | $\Phi_{i,j}$ |
|---|---|---|---|---|
| Self | 0.00000 | 0.00000 | 1.00000 | 0.50000 |
| Parent-offspring pair | 0.00000 | 1.00000 | 0.00000 | 0.25000 |
| Full siblings | 0.25000 | 0.50000 | 0.25000 | 0.25000 |
| Second degree | 0.50000 | 0.50000 | 0.00000 | 0.12500 |
| Third degree | 0.75000 | 0.25000 | 0.00000 | 0.06250 |
| Fourth degree | 0.87500 | 0.12500 | 0.00000 | 0.03125 |
| Unrelated | 1.00000 | 0.00000 | 0.00000 | 0.00000 |

**Table S2.** Summary statistics of execution time (in seconds) of 100 simulations of the genealogy scenario 1 using different network methods implemented in NAToRA. The scenario 1 consists of four generations with 10 male and 10 female in each one of the generations (total 80 individuals) and a proportion of unrelated individuals of 0.5.

| **Scenario 1** | **Minimum** | **Average** | **Standard deviation** | **Maximum** |
|---|---|---|---|---|
| **Node Degree Centrality** | 0.005 | 0.011 | 0.003 | 0.028 |
| **Node Betweenness Centrality** | 0.023 | 0.059 | 0.024 | 0.127 |
| **Closeness Centrality** | 0.013 | 0.026 | 0.009 | 0.059 |
| **Optimal Result (Clique)** | 0.007 | 0.019 | 0.024 | 0.193 |

**Table S3 . S**ummary statistics of execution time (in seconds) of 100 simulations of the genealogy scenario 2 using different network methods implemented in NAToRA. The scenario 2 consists of four generations with 50 male and 50 female in each of the 4 generations (total 400 individuals) with proportion of unrelated of 0.5.

| **Scenario 2** | **Minimum** | **Average** | **Standard deviation** | **Maximum** |
|---|---|---|---|---|

| | | | | |
|---|---|---|---|---|
| **Node Degree Centrality** | 0.064 | 0.092 | 0.021 | 0.154 |
| **Node Betweenness Centrality** | 1.314 | 2.341 | 0.484 | 3.498 |
| **Closeness Centrality** | 0.294 | 0.471 | 0.106 | 0.787 |
| **Optimal Result (Clique)** | 0.045 | 15,585.9 | 154,119.8 | 1,541,290.4 |

**Table S4.** Summary statistics of execution time (in seconds) of 98 simulations of the genealogy scenario 3 using different network methods implemented in NAToRA. In this scenario we simulated 100 times a scenario with 100 male and 100 female in each of the 4 generations (total 800 individuals) with proportion of unrelated of 0.5. This table has the results for 98 of the 100 tests. The remaining tests are not available due to the large computational time (> 7 months).

| Scenario 3 | Minimum | Average | Standard deviation | Maximum |
|---|---|---|---|---|
| **Node Degree Centrality** | 0.216 | 0.265 | 0.043 | 0.436 |
| **Node Betweenness Centrality** | 11.010 | 15.690 | 1.961 | 20.400 |
| **Closeness Centrality** | 1.256 | 1.747 | 0.270 | 2.624 |
| **Optimal Result (Clique)** | 0.106 | 5,968.8 | 52,819.9 | 522,183.7 |

**Table S5.** Summary statistics of execution time (in seconds) of 87 simulations of the genealogy scenario 4 using different network methods implemented in NAToRA. In this scenario we simulated 100 times a scenario with 500 males and 500 females in each of the 4 generations (total 4000 individuals) with proportion of unrelated of 0.5. This table has the results for 87 of the 100 tests. The remaining tests are not available because of the large computational time (> 7 months).

| Scenario 4 | Minimum | Average | Standard deviation | Maximum |
|---|---|---|---|---|
| **Node Degree Centrality** | 3.526 | 4.907 | 0.915 | 8.415 |
| **Node Betweenness Centrality** | 3,190.0 | 3,877.0 | 484.11 | 5,565.0 |

| | | | | |
|---|---|---|---|---|
| **Closeness Centrality** | 33.840 | 46.110 | 8.126 | 71.650 |
| **Optimal Result (Clique)** | 1.300 | 87,204.2 | 253,790.6 | 1,409,964.3 |

**Table S6. Experimental results with real datasets.** We tested (1) three different datasets in relation to (2) the number of individuals pruned and (3) the execution time of four network methods implemented in NAToRA.

| | | **BAMBUÍ** | **SHIMAA** | **GUZERÁ** |
|---|---|---|---|---|
| **1 - Description of Databases** | Number of individuals | 1,442 | 42 | 1,036 |
| | Number of related individuals | 885 | 32 | 1021 |
| **2 - Number of individuals pruned by method** | Node Degree Centrality | 522 | 18 | 828 |
| | Node Betweenness Centrality | 525 | 18 | 849 |
| | Closeness Centrality | 527 | 18 | 845 |
| | Optimal Result (Clique) | 520 | 18 | -* |
| **3 - Runtime by method (in seconds)** | Node Degree Centrality | 1.227 | 0.031 | 17.2 |
| | Node Betweenness Centrality | 383.715 | 0.100 | 6,086.2 |
| | Closeness Centrality | 11.705 | 0.055 | 1,484.4 |
| | Optimal Result (Clique) | 2.632 | 0.037 | -* |

*The results of the execution of the optimal algorithm are not available for Guzerá because of the large computational time (> 7 months)

**Table S7. Comparison of relatedness-pruning results to generate datasets with only kinship relationships below second-degree. Methods are PLINK (--rel-cutoff), KING (--degree 2 --unrelated), NAToRA and the general strategy (kinship>2nd).** Cutoff values are 0.1768 for the relationship coefficiente (calculated by PLINK) and degree 2 for the kinship coefficient (calculated by KING). Because this was the value used by --degree 2 according to the tool manual. Values in the Original Dataset Size and Unrelated Dataset Size columns are in the format: number of individuals on the dataset (number of relationships on the dataset). For NAToRA we show the heuristic result. The time column is the system time of each execution. The times of NAToRA and kinship < 2nd degree are smaller because the relationship was calculated previously, different from KING and PLINK that calculate the relationship at the same run of pruning process.

| Database | Relatedness estimated by | Original Dataset Size | Parameters to exclude by method | Unrelated Dataset Size | Time |
|---|---|---|---|---|---|
| BAMBUI | PLINK | 1442(1572) | plink --rel-cutoff 0.1768 | 908(234) | 7m15.915s |
| | | | kinship < 2nd degree | 491(0) | 0m6.773s |
| | | | NAToRA -c 0.1768 | 869(0) | 0m8.912s |
| | KING | 1442(920) | king --degree 2 --unrelated | 839 (1) | 0m53.866s |
| | | | kinship < 2nd degree | 602(0) | 0m4.527s |
| | | | NAToRA --degree 2 | 949(0) | 0m8.719s |
| SHIMAA | PLINK | 45(95) | plink --rel-cutoff 0.1768 | 26(8) | 0m0.355s |
| | | | kinship < 2nd degree | 10(0) | 0m0.037s |
| | | | NAToRA -c 0.1768 | 23(0) | 0m1.573s |
| | KING | 45(68) | king --degree 2 --unrelated | 45(68) | 0m14.989s |
| | | | kinship < 2nd degree | 12(0) | 0m0.036s |
| | | | NAToRA --degree 2 | 25 (0) | 0m1.828s |
| GUZERA | PLINK | 1036(17875) | plink --rel-cutoff 0.1768 | 210(369) | 0m7.165s |
| | | | kinship < 2nd degree | 3(0) | 0m3.618s |
| | | | NAToRA -c 0.1768 | 176(0) | 0m7.339s |
| | KING | 1036(12861) | king --degree 2 --unrelated | 88(0) | 0m16.256s |

| | | | | | |
|---|---|---|---|---|---|
| | | | kinship < 2nd degree | 24(0) | 0m2.355s |
| | | | NAToRA --degree 2 | 218(0) | 0m7.255s |
| EPIGEN | PLINK | 6487(2965) | plink --rel-cutoff 0.1768 | 5600 (880) | 321m10.367s |
| | | | kinship < 2nd degree | 4446(0) | 0m6.048s |
| | | | NAToRA --degree 2 | 5355(0) | 2m55.384s |
| | KING | 6847 (1090) | king --degree 2 --unrelated | 5691(7) | 2m18.752s |
| | | | kinship < 2nd degree | 5321(0) | 0m6.395s |
| | | | NAToRA --degree 2 | 5832(0) | 2m13.103s |

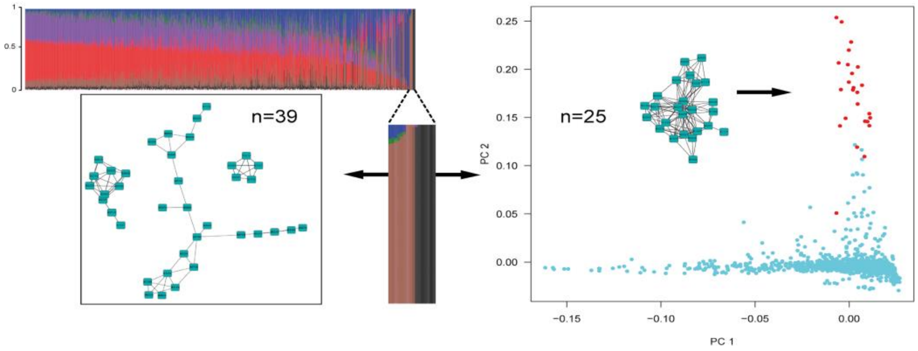


**Figure S1. Familiar structure in Bambuí identified by ADMIXTURE in K=7.** Using Unsupervised ADMIXTURE analysis for BAMBUI dataset the algorithm identifies two ancestry clusters (brown and black) that match with a set of related individuals estimated by REAP. Individuals from black component were identified by the second component of PCA (red dots). Figure from Kehdy *et al*. 2015.

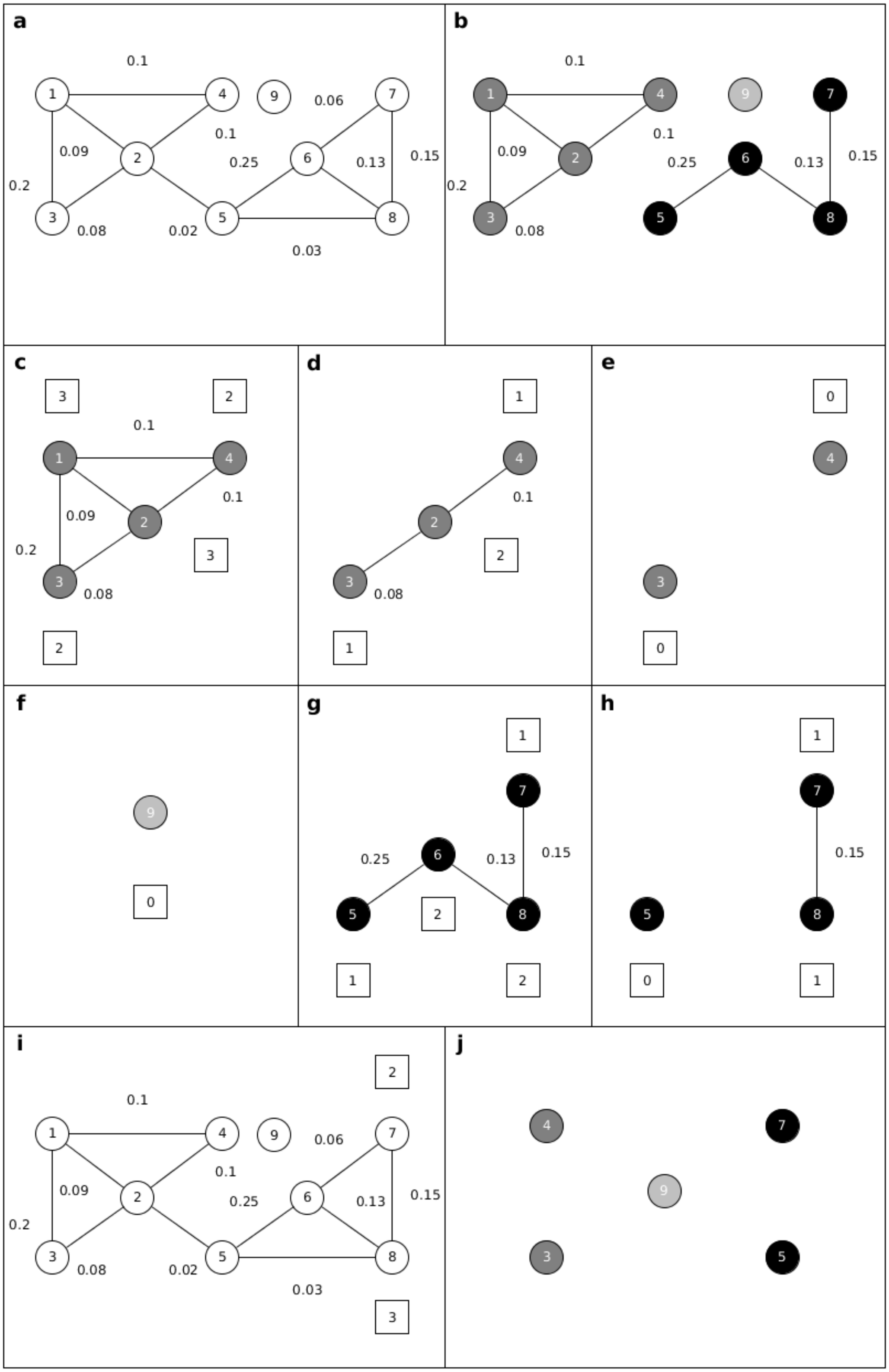
a
0.1
1
4
9
0.06
7
0.1
0.09
2
0.25
6
0.13
0.15
0.2
3
0.08
0.02
5
8
0.03
b
0.1
1
4
9
7
0.1
0.09
2
0.25
6
0.13
0.15
0.2
3
0.08
5
8
c
3
2
0.1
1
4
0.1
0.09
2
3
0.2
3
0.08
2
d
1
4
0.1
2
2
3
0.08
1
e
0
4
3
0
f
9
0
g
1
7
0.15
0.25
6
0.13
5
2
8
1
2
h
1
7
0.15
5
8
0
1
i
2
0.1
1
4
9
0.06
7
0.1
0.09
2
0.25
6
0.13
0.15
0.2
3
0.08
0.02
5
8
0.03
3
j
4
7
9
3
5

**Figure S2.** NAToRA Algorithm example. In (a) is represented the network with all kinship coefficient values (network N). If we were to remove all related individuals from network N, only individual 9 would remain. In (b) is represented the network $N_c$ (Network with cuts) using cutoff value 0.07. After generating $N_c$ the algorithm detects the network families, represented as individuals with the same node color. In (c),(d) and (e) we show the node elimination process for the dark-grey colour family. The process consists of calculating the centrality metric (in the figure, we used node degree centrality; numbers in white boxes are the individuals' node degree centrality) and eliminating the individual with highest centrality metrics until only nodes with at most one edge remain. In (c), there was a centrality tie between individuals 1 and 2 and we eliminate individual 1, because it had heavier edges. In (f) the light-grey colour family was analysed. In (g), (h) and (i), we illustrate the node elimination process for the black colour family. In (g) we see that individual 6 is eliminated because it has a high centrality degree and largest sum of edges' weights. (h) shows a case where elimination based on centrality metrics is inefficient, since both connected individuals have the same value. In (i) we show a case of tie breaker, where the algorithm gets the original database (presented in (a)), selects the edges within the range selected by the user (in the example values are 0.03 and 0.20), calculates the centrality and excludes the individual with highest centrality. The output is presented in (j) with a list of individuals that should remain in the dataset.

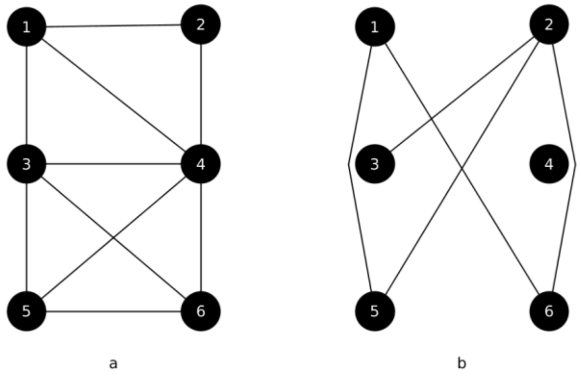


**Figure S3.** Example of a network (a) and its complementary network (b). There is an edge in (b) if and only if it does not exist in (a).

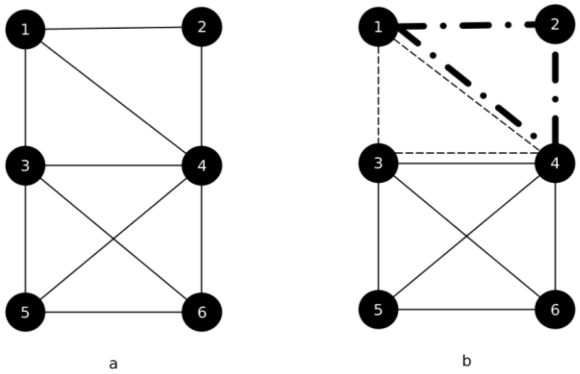


**Figure S4.** Example of cliques on a network. In (a) we have a sample network and in (b) we show all maximal cliques represented by the shape of edges. A maximal clique is a clique that cannot be increased by adding more adjacent nodes, i.e., a clique that does not belong to a larger clique. The set {3,4,5,6} is the maximum clique, that is, the biggest maximal clique in the network.

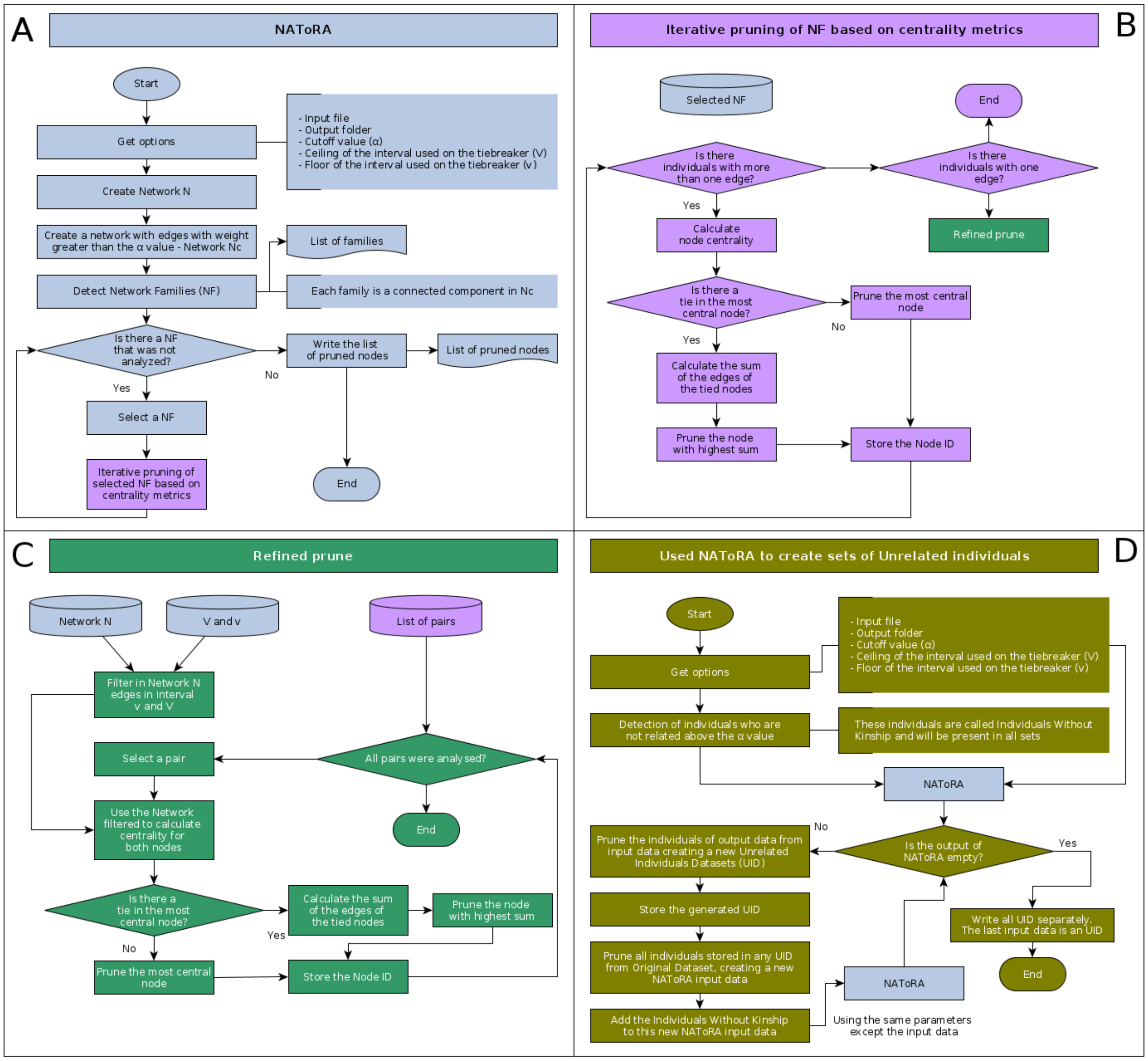


**Figure S5. NAToRA workflow.** A) Overview of the NAToRA algorithm; B) Iterative pruning of the Network Families (NF) based on centrality metrics; C) Refined pruning; and D) The using of NAToRA to create unrelated set of individuals. The colors were used to facilitate the differentiation of the steps as well as to clarify the origin of the input of each step.

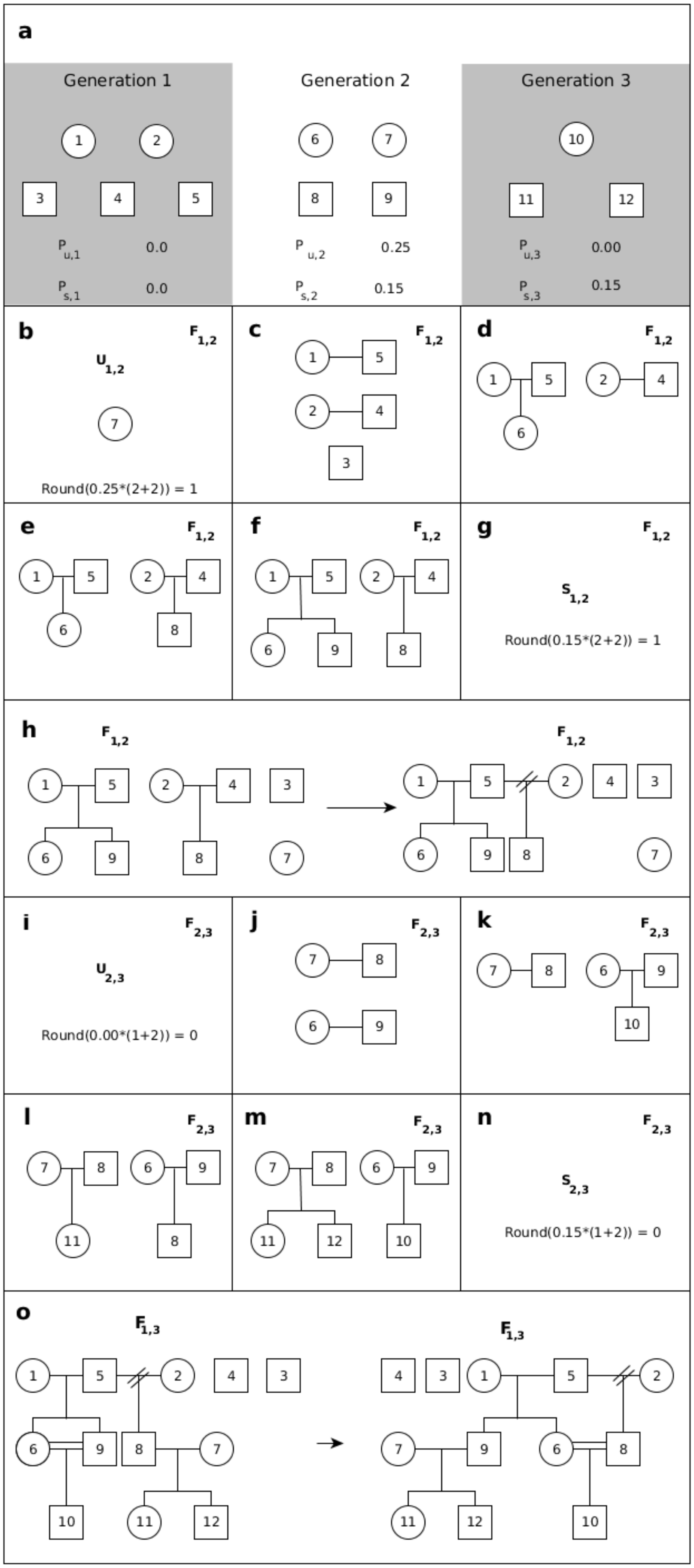
a
Generation 1
Generation 2
Generation 3
P u,1 0.0
P s,1 0.0
P u,2 0.25
P s,2 0.15
P u,3 0.00
P s,3 0.15
b
U 1,2
F 1,2
Round(0.25*(2+2)) = 1
c
F 1,2
d
F 1,2
e
F 1,2
f
F 1,2
g
F 1,2
S 1,2
Round(0.15*(2+2)) = 1
h
F 1,2
F 1,2
i
F 2,3
U 2,3
Round(0.00*(1+2)) = 0
j
F 2,3
k
F 2,3
l
F 2,3
m
F 2,3
n
F 2,3
S 2,3
Round(0.15*(1+2)) = 0
o
F 1,3
F 1,3

**Figure S6. Genealogy simulator example.** In (a) we represented the input data (generation 1, generation 2 and generation 3), where circles represent female, square represent male, the values of $P_{u,g}$ and $P_{s,g}$ for each generation (0.0 and 0.0 for g=1; 0.25 and 0.15 for g=2; 0.0 and 0.15 for g=3). In (b) we show the calculation $(A_{u,2})$ and selection of the set of unrelated individuals($U_2$) for the sub-heredogram $F_{1,2}$. In (c) we have the formation of all couples of generation 1 and in (d), (e) and (f) the distribution of related individuals for these couples. In (g) we show the calculation $(A_{s,2})$ of the minimum number of step brothers to $F_{1,2}$. As the value was greater than 0 in (h) have the generation of half-brothers by replacing a parent of a couple randomly selected. After this the algorithm completes the generation of sub-heredogram $F_{1,2}$ and in steps (i), (j), (k), (l), (m) and (n) the steps for $F_{2,3}$. Having generated all sub-heredograms, the algorithm starts the last step which is join all sub-heredogram to generate a single heredogram. The algorithm begins by mapping the children of $F_{1,2}$ as the parents of $F_{2,3}$, making the joining. After joining, the algorithm looks for inconsistencies, that is, looks for siblings who have children together. This problem is represented in (o) where 6 and 9 (siblings $F_{1,2}$) in have a son in $F_{2,3}$. To solve the algorithm randomly select one of the two siblings of generation $F_{1,2}$ and replace it with another individual with other same-sex and generation individual and search again for inconsistencies. As the process involves steps with randomness the same scenario can generate several different genealogies.

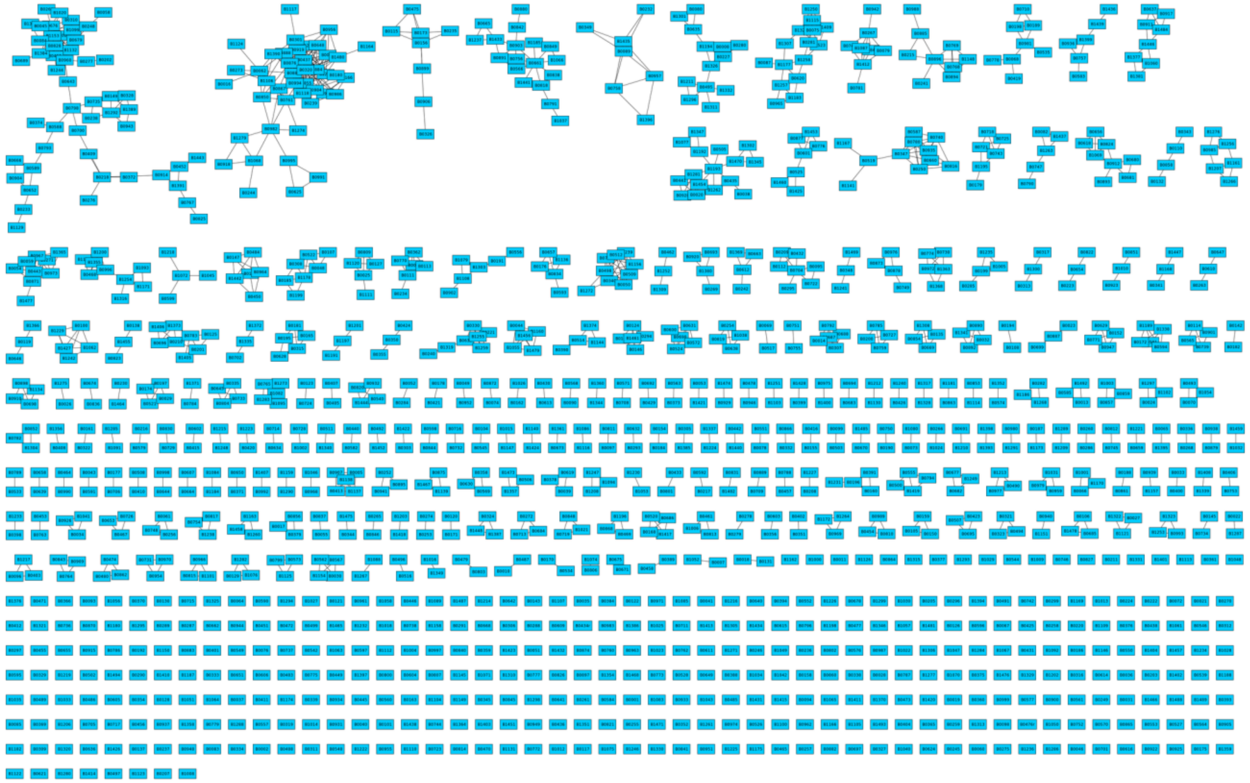

**Figure S7. Bambuí (BAMBUI) kinship network.** We inferred the kinship coefficient using REAP and considered related individuals with kinship coefficient greater than 0.1.

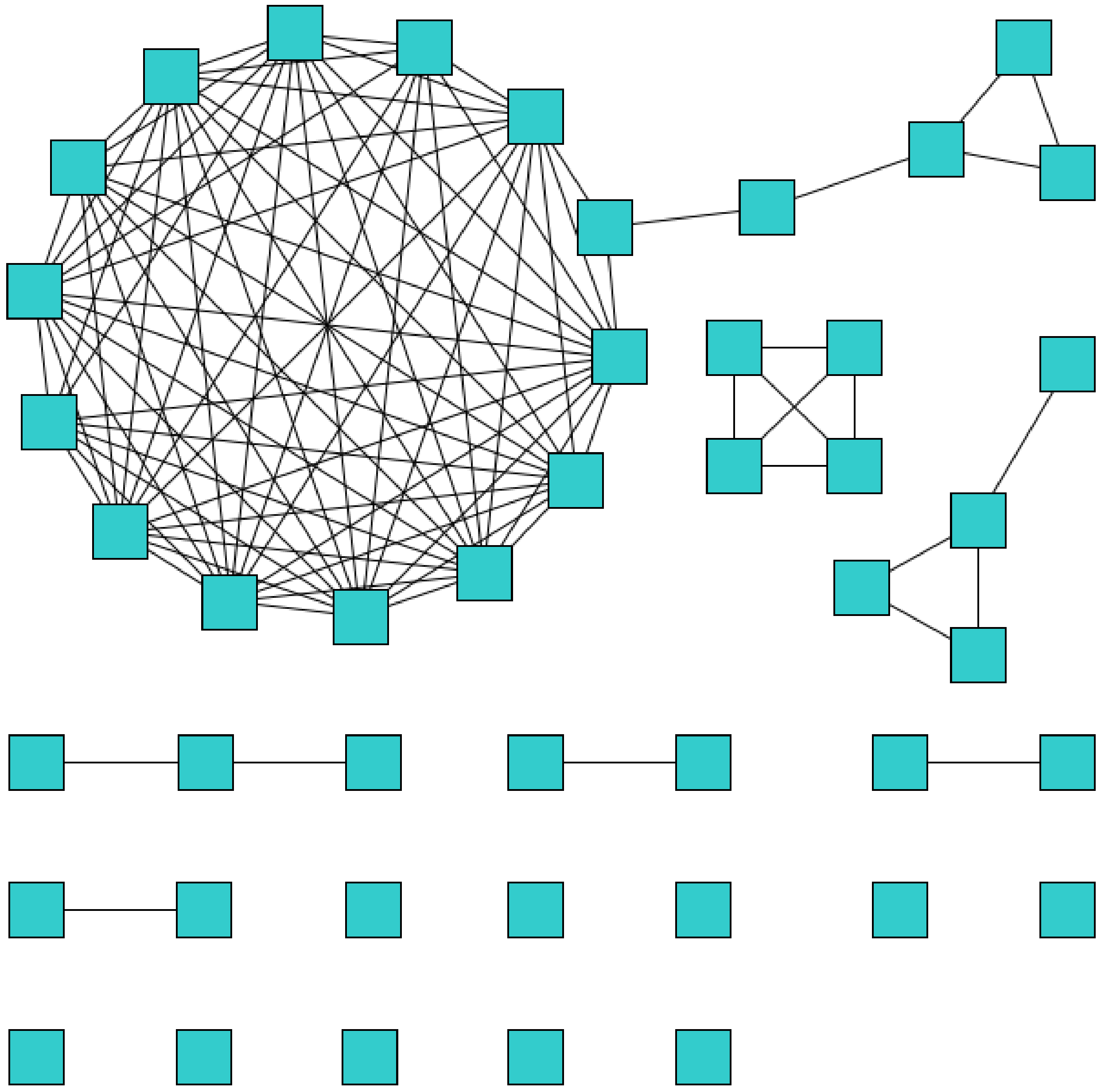

**Figure S8. Matsiguenkas from Amazon Yunga (SHIMAA) kinship network.** We inferred the kinship coefficient using PLINK and considered related individuals with kinship coefficient greater than 0.0884.

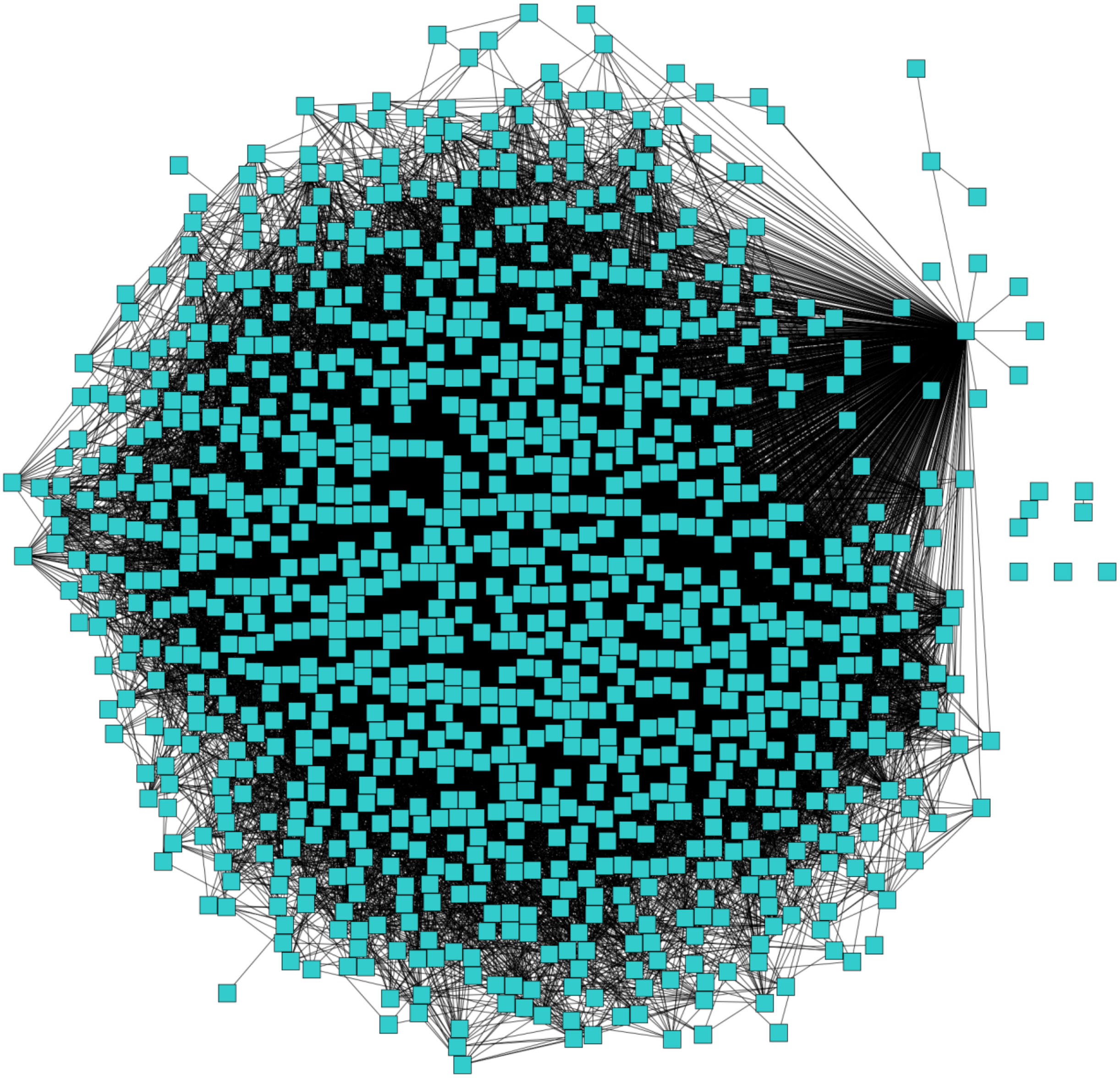

**Figure S9. National Program for Improvement of Guzera for Milk (GUZERA) kinship network.** We inferred the kinship coefficient using PLINK and considered related individuals with kinship coefficient greater than 0.1.

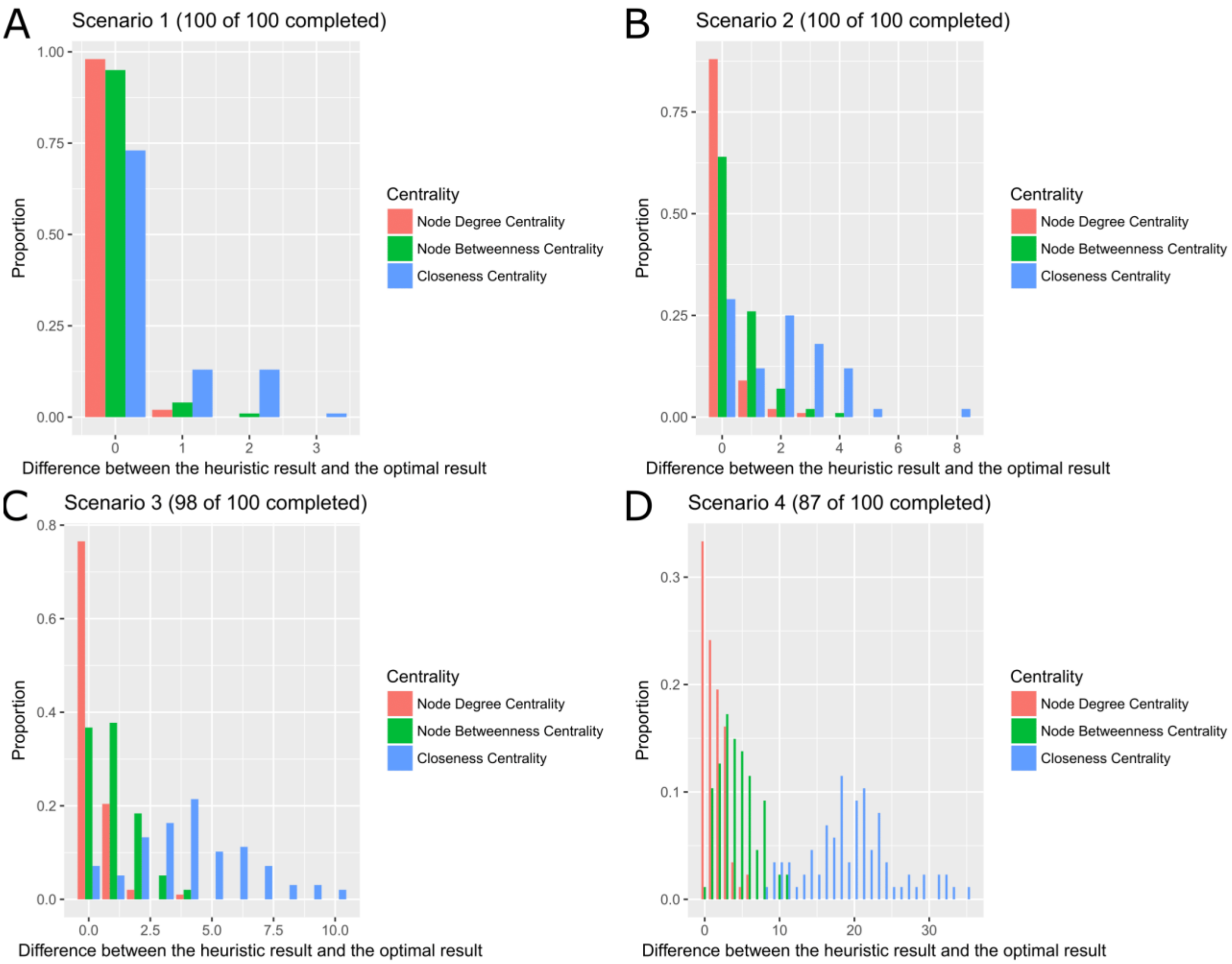


**Figure S10. Comparison between heuristics and optimal.** Distribution of the difference between heuristic and optimal results for all scenarios. In (A) the results for Scenario 1, in which the node degree centrality and node betweenness centrality obtained almost all results similar to the optimal algorithm. In (B) Scenario 2 the difference starts to increase between node degree centrality and node betweenness, which is widened in Scenarios 3 (C) and 4 (D).

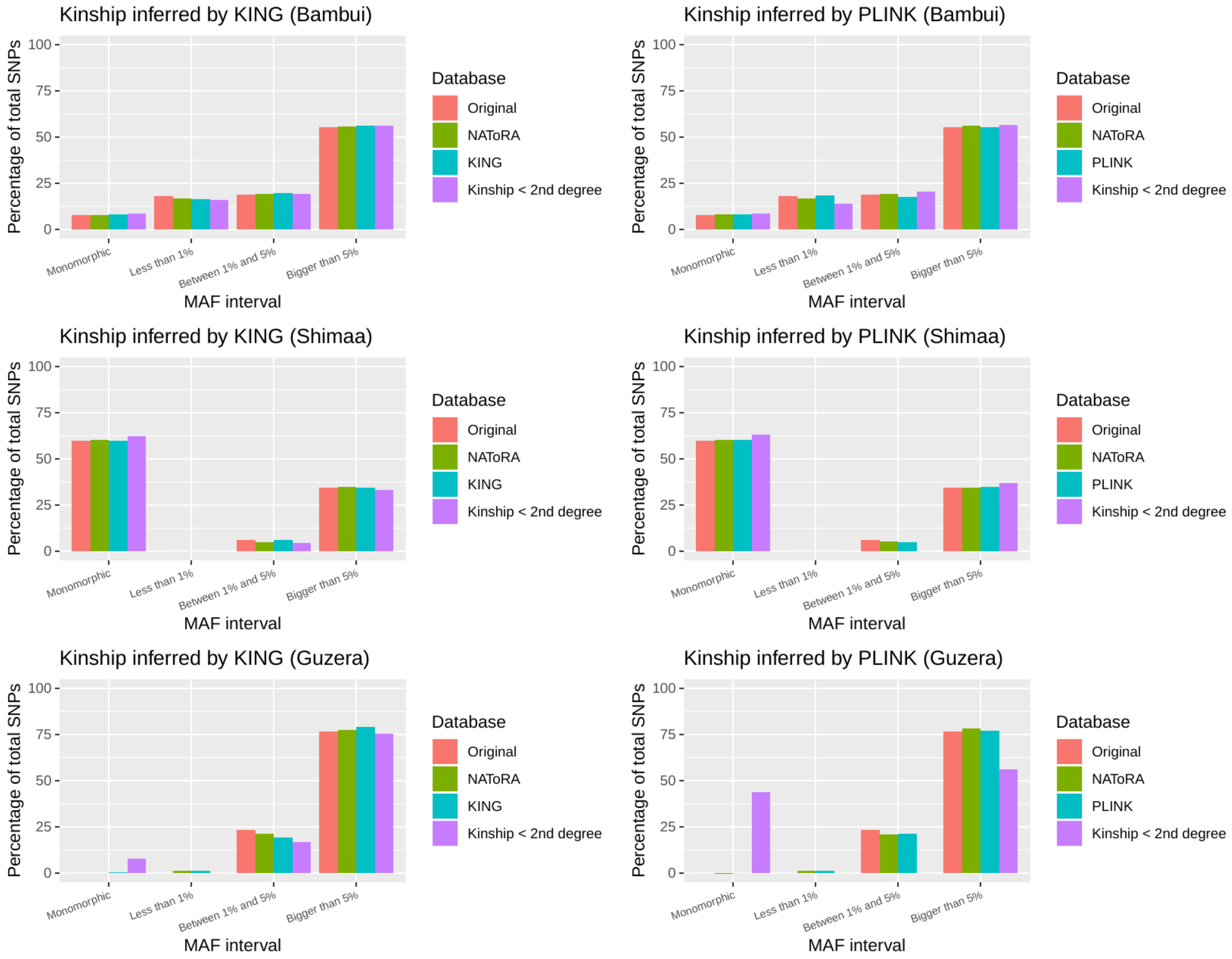


**Figure S11. Comparison of the percentage of total SNPs in the dataset present in each MAF interval before (Original) and after different relatedness-pruning methods (PLINK or KING, NAToRA and Kinship < 2nd degree).** We divided the SNPs into four categories: Ultra rare (0< MAF ≤ 0.01), rare (0.01 < MAF ≤ 5%), common (MAF > 5%) and monomorphics (MAF = 0).

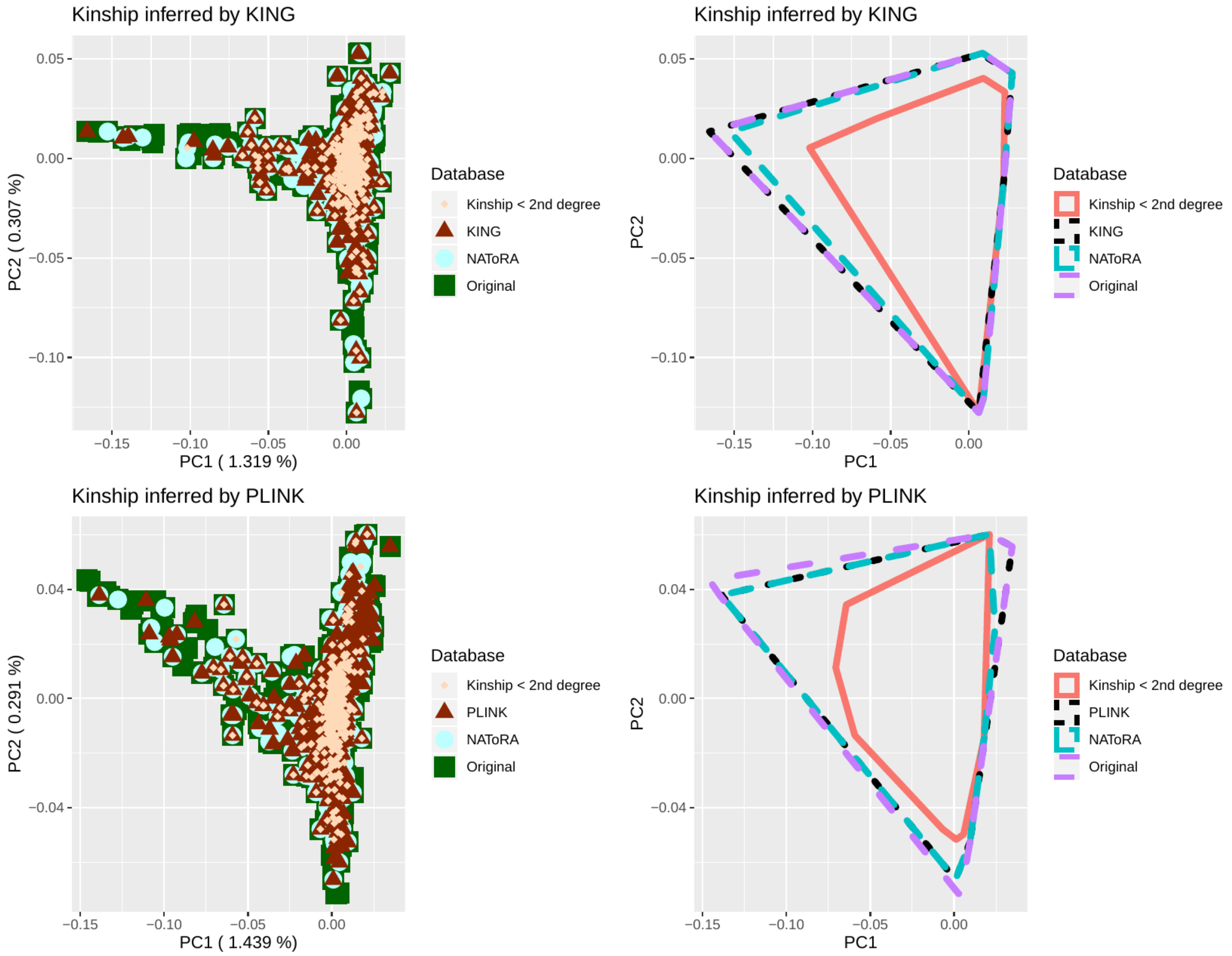


**Figure S12. Principal Component Analysis for BAMBUÍ dataset and the respective resampling (tool removing, NAToRA and Kinship < 2nd degree) performed together.** The figure is composed of 2 components and with it we show the representativeness of resampling techniques in the original base (the polygon in B and D is intended to facilitate visualization showing the area covered by the resampling technique). The values within the parentheses are the percentages of variance explained by the PC

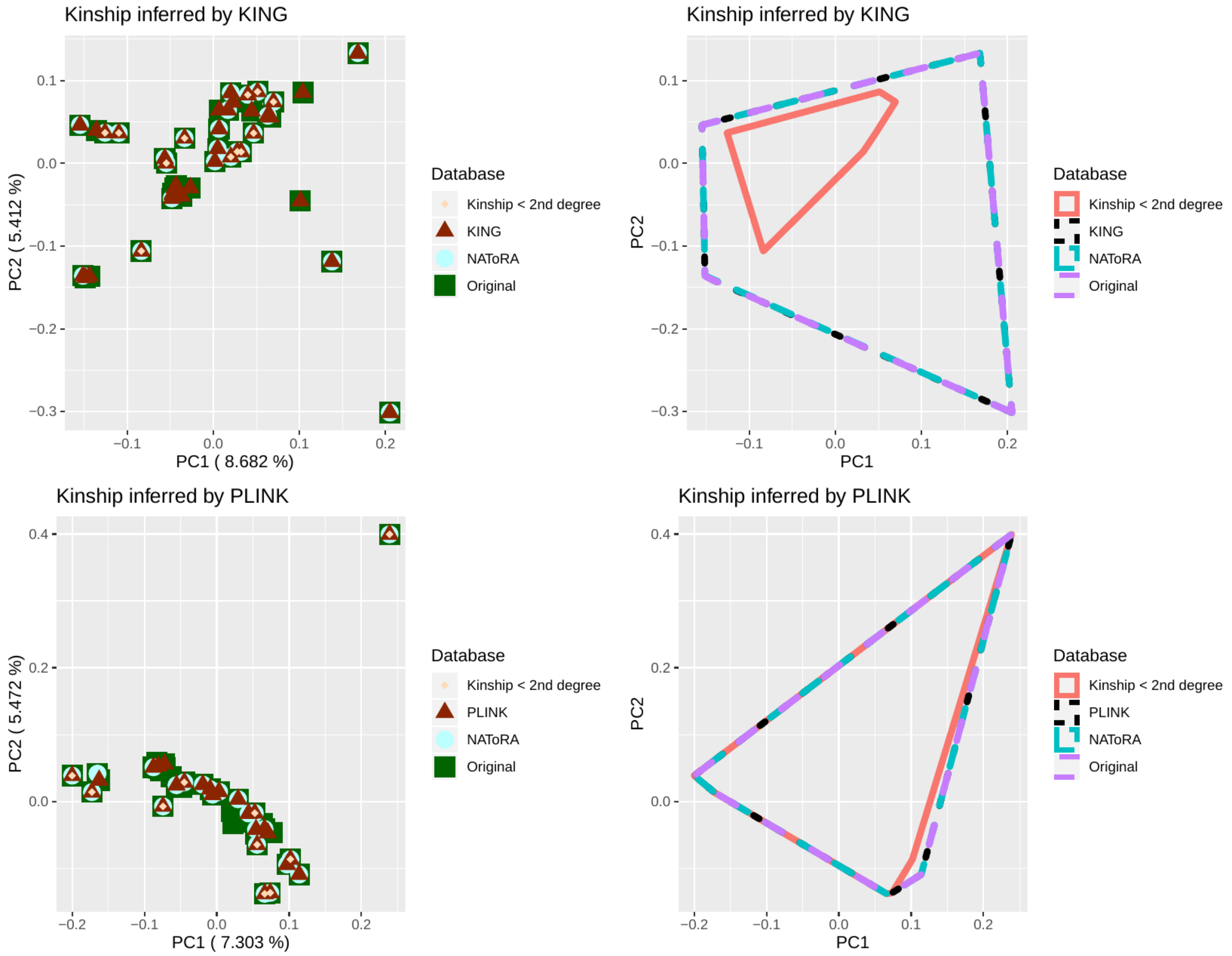


**Figure S13. Principal Component Analysis for SHIMAA dataset and the respective resampling (tool removing, NAToRA and Kinship < 2nd degree) performed together.** The figure is composed of 2 components and with it we show the representativeness of resampling techniques in the original base (the polygon in B and D is intended to facilitate visualization showing the area covered by the resampling technique). The values within the parentheses are the percentages of variance explained by the PC

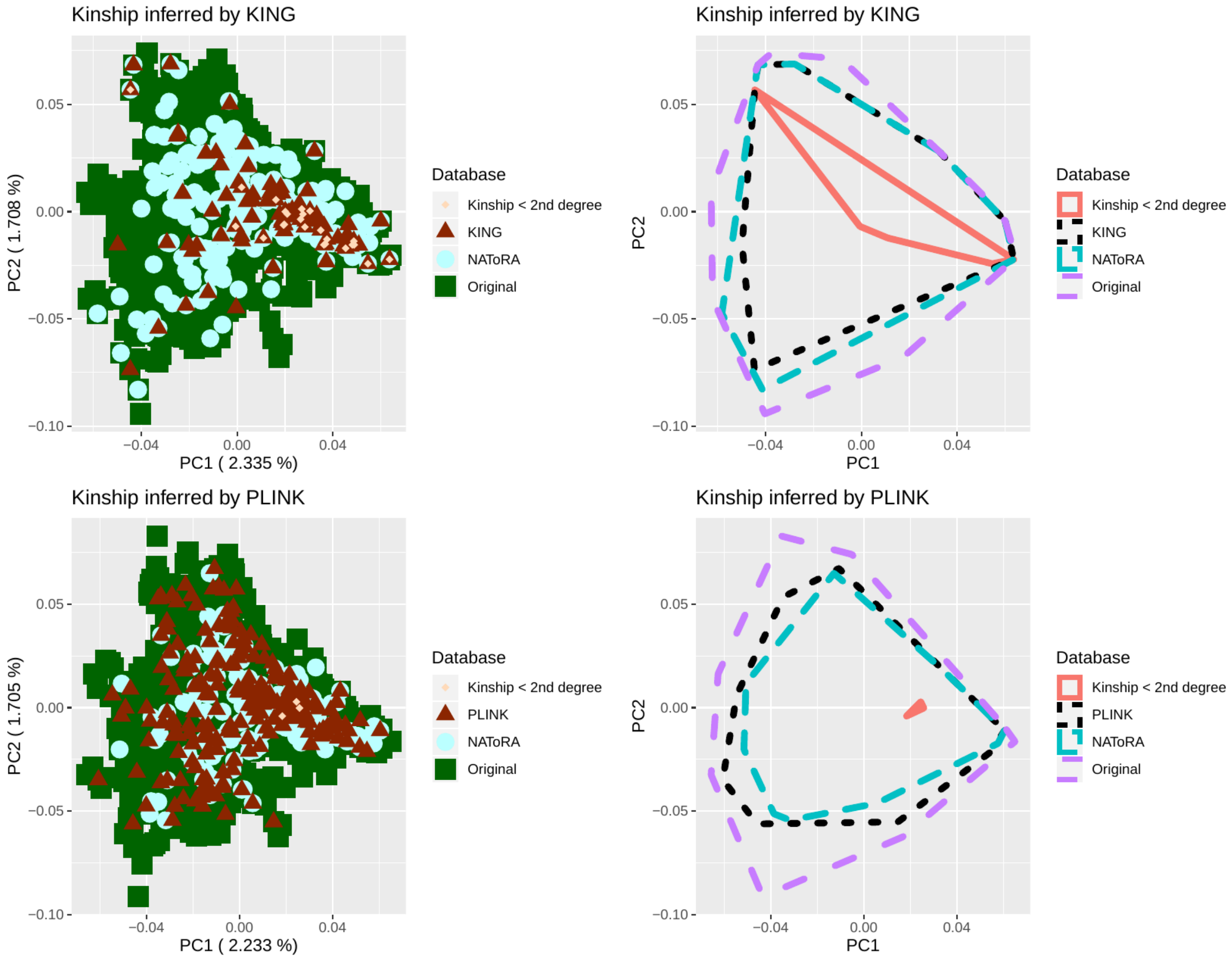


**Figure S14.Principal Component Analysis for Guzerá dataset and the respective resampling (tool removing, NAToRA and Kinship < 2nd degree) performed together.** The figure is composed of 2 components and with it we show the representativeness of resampling techniques in the original base (the polygon in B and D is intended to facilitate visualization showing the area covered by the resampling technique). The values within the parentheses are the percentages of variance explained by the PC.

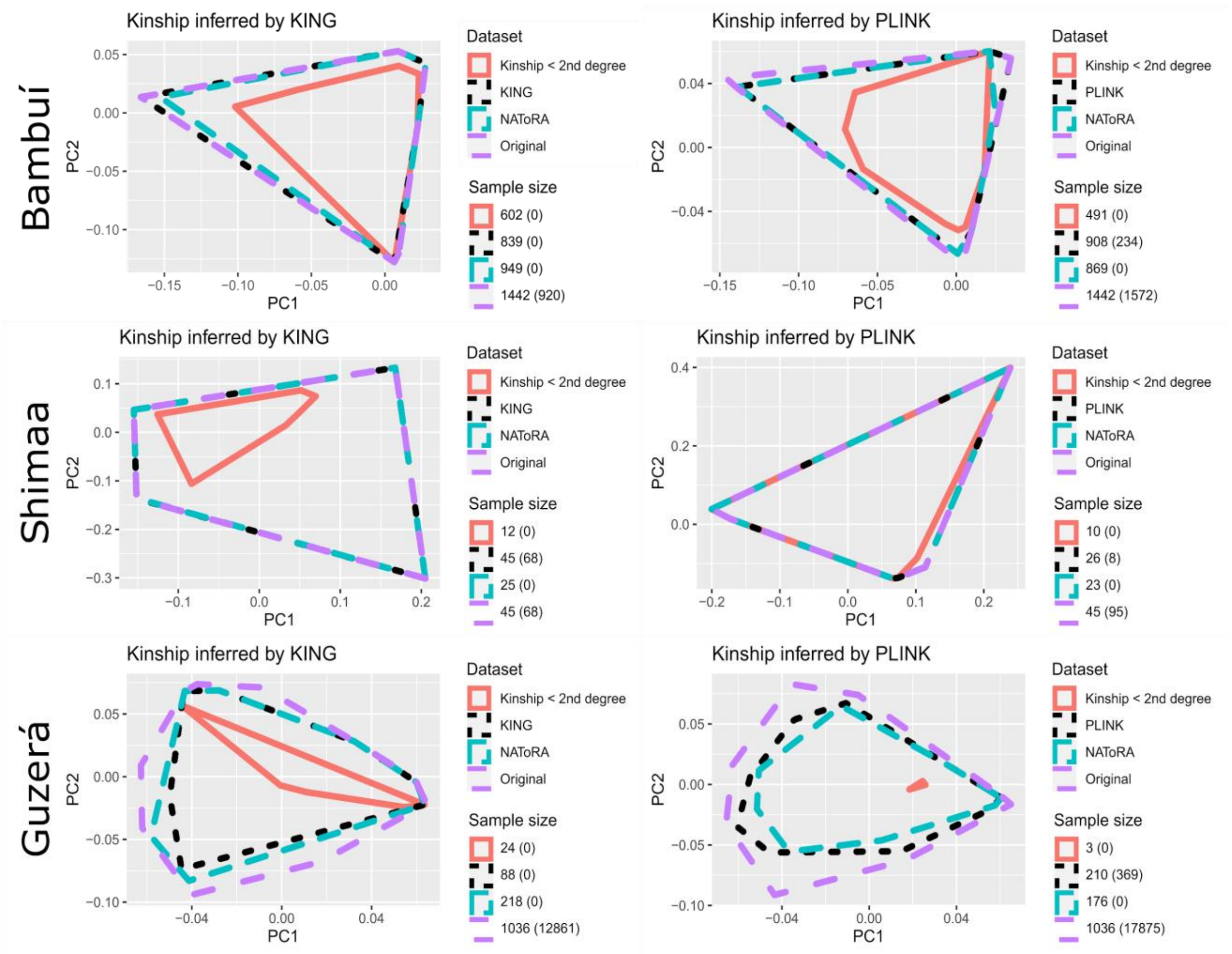


**Figure S15. Principal Component Analysis for all datasets and the respective resampling (PLINK or REAP, NAToRA and Kinship < 2nd degree) performed together.** The figure is composed of 2 components and with it we show the representativeness of resampling techniques in the original base. The number in the legend is the number of samples and number of relationships (inside the brackets) for all subsets.

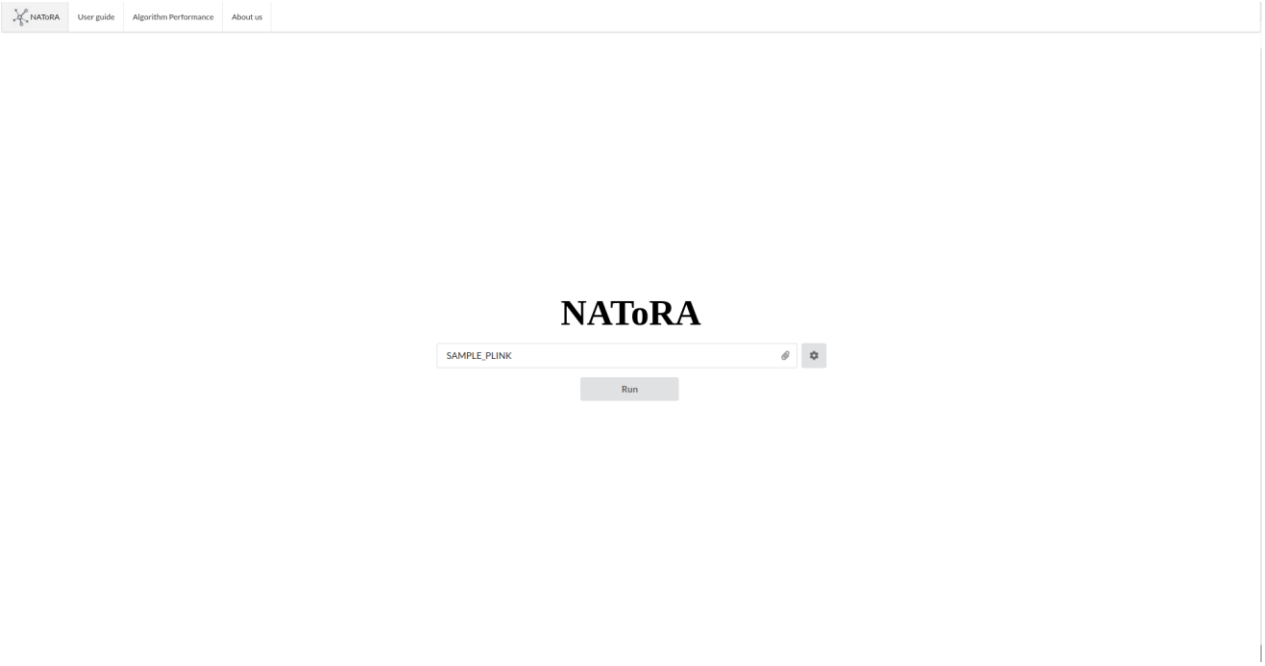


**Figure S16. NAToRA’s webtool main page.**

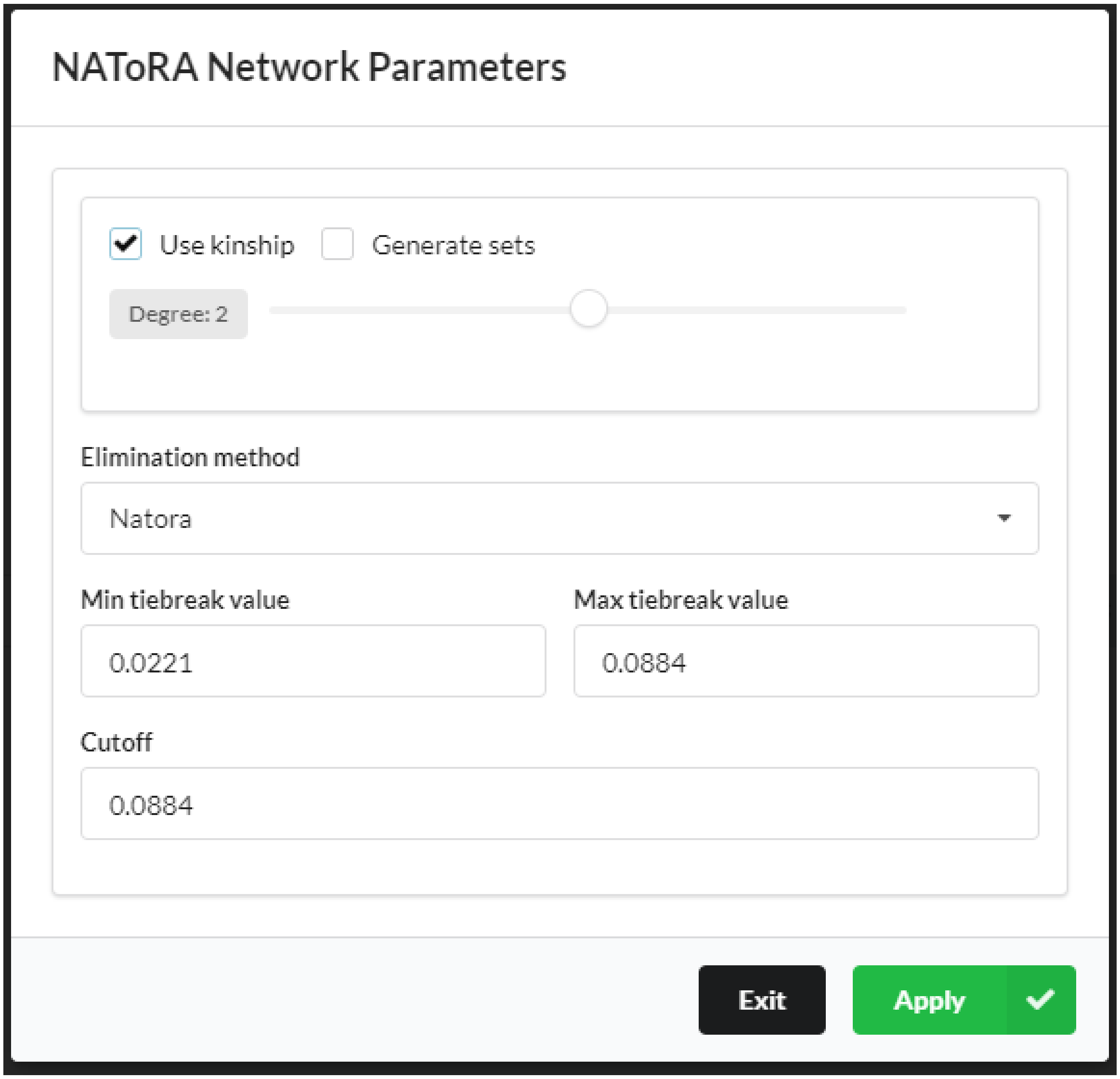


**Figure S17. NAToRA network parameters menu.** You can access this menu by clicking on the gear button. You can define the relatedness cutoff in two different ways: (i) if your relatedness metric is Kinship coefficient you can select the "Use kinship" that will set the default values in Min and Max tiebreaker and Cutoff or (ii) manually setting these values typing in the respective boxes. If your relatedness metric is different from Kinship Coefficient, such as PI_HAT, you should use the second option. In this example we selected "Use kinship" and 2$^{nd}$ degree, which configures the values of cutoff (0.0884), min and max tiebreaker values.

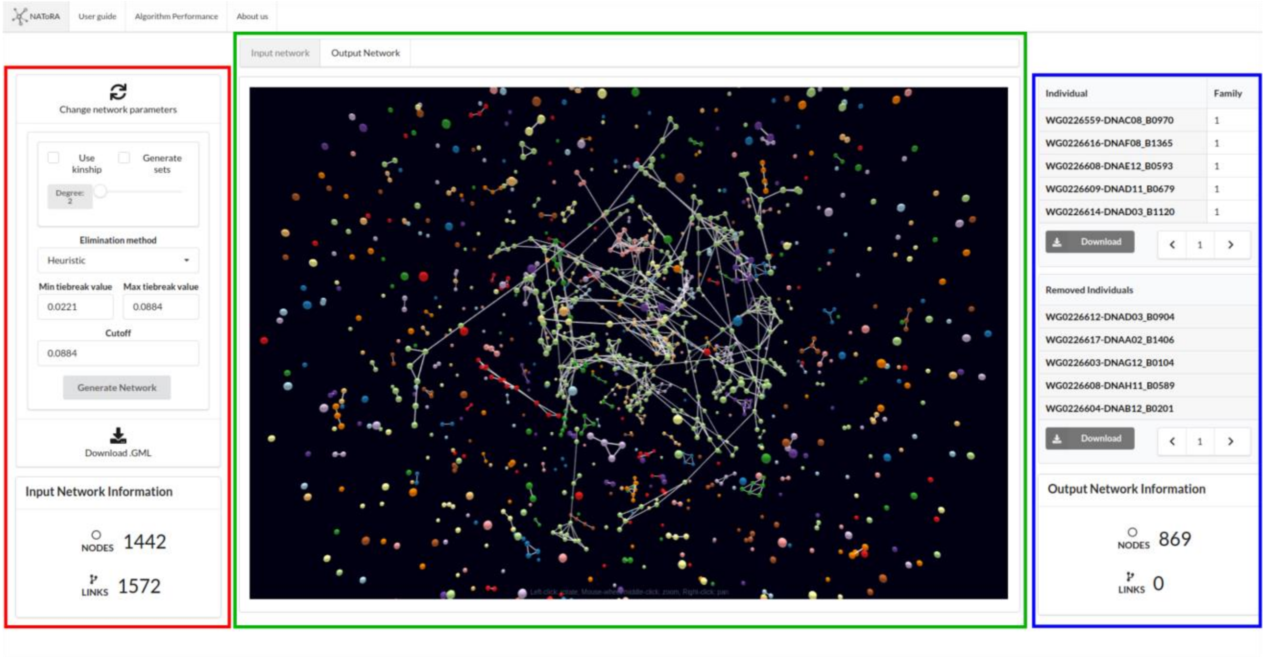


**Figure S18. NAToRA output screen.** This screen was generated by the input file (SAMPLE_PLINK) and the parameters that were chosen. At the left (red) we have the input information, with the menu to re-run the analysis without uploading the file again, and the input network file. You can also download the GML file of your network with the cutoff value. In the middle (green) we present the network view that allows the user to visualize the input network with the kinship cutoff and the output network (without relationships greater than the selected cutoff). In the right (blue) is the output results, that you can download the family list file and individuals to be removed from your data to get a data without relationship greater than the cutoff value. You can also see the number of individuals and relationships left.

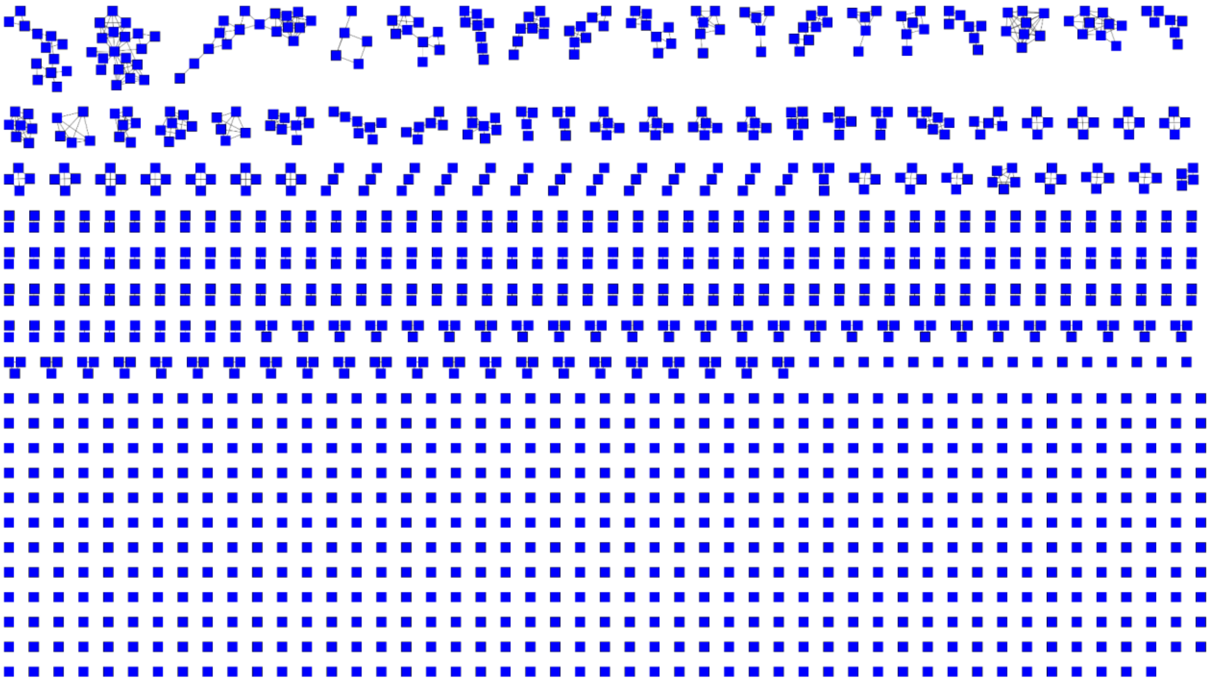

**Figure S19. The BAMBUI network with kinship inferred by KING.** After the inference we used the script KING2NAToRA.pl to create the file with the format of NAToRA input file. With the file with NAToRA format we used the generateGML.pl to convert to GML format. We used as cutoff value 0.0884 (second degree) to create this plot.

A B

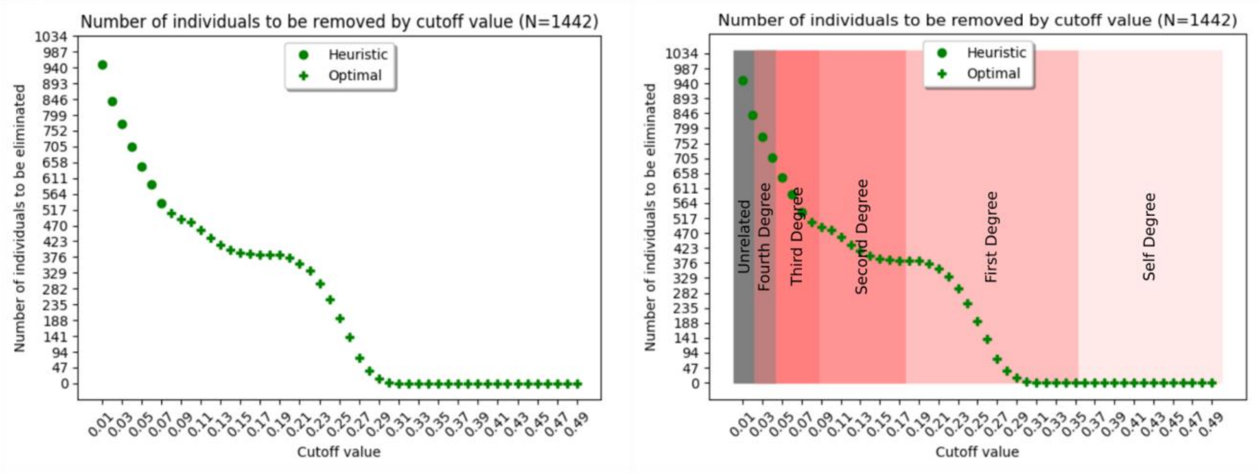


C

```
Values by degree                  Min                          Theoretical    Max
Self degree interval:    0.3535533905932738                0.5             0.5
First degree interval:   0.1767766952966369                0.25            0.3535533905932738
Second degree interval:  0.08838834764831845               0.125           0.1767766952966369
Third degree interval:   0.04419417382415922               0.0625          0.08838834764831845
Fourth degree interval:  0.02209708691207961               0.03125         0.04419417382415922
```

**Figure S20. An example of the output of "--test 0.5" and "--test 0.5 --kinship" parameters applied to BAMBUI dataset with kinship coefficient calculated by KING.** (A) Result for the "--test" parameter, in which NAToRA calculates the estimated sample loss for 50 cutoff values (beginning on 0.05 to the max value informed by the user, in this example 0.5). Sample loss is shown for the heuristic and optimal (when possible) versions of the algorithm. (B) Same result shown in (A) with added colours to inspect the intervals of each kinship value used by KING. (C) Kinship values for each kinship degree used by KING presented by the flags "--test --kinship". If the user wants to remove second degree and above, the value to -c flag is 0.0884, for example.

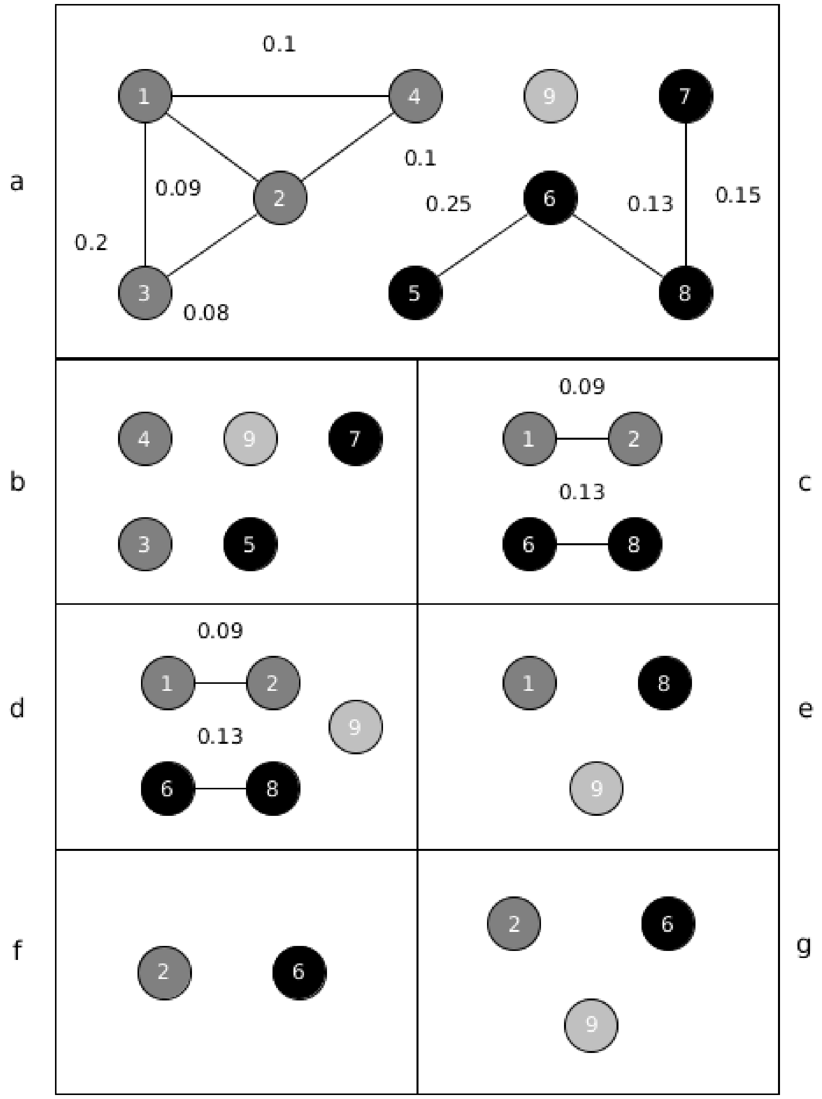


**Figure S21 . Example of using NAToRA to generate independent data sets.** In (a) we show a $N_c$network. The Unrelated Dataset (DU) is DU={9}. In (b) shows the Analysis Dataset 1 ($AD_1$), composed by $AD_1$={3,4,5,7,9}. In (c) is presented the network $N_c$without individuals present in $AD_1$. In (d) we insert the individuals present in DU and we execute NAToRA in this dataset, (e) generating the $AD_2$={1,8,9}. In (f) we remove from dataset generated in (d) the individuals from $AD_2$. In (g) we insert the DU dataset and run NAToRA. Since output from NAToRA is empty, this means there is no relationship on this network, so $AD_3$={2,6,9}.

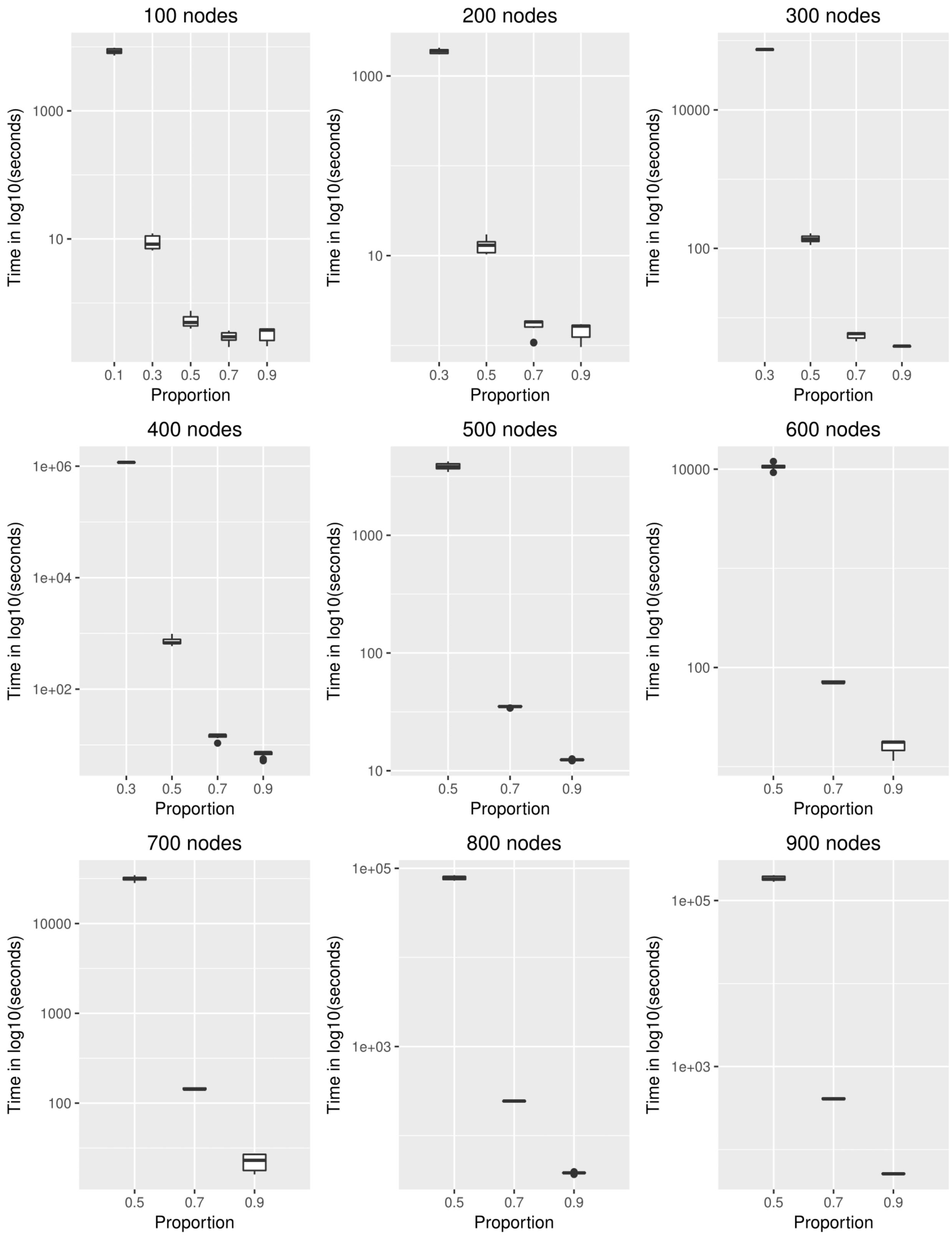
100 nodes
Time in log10(seconds)
1000
10
0.1
0.3
0.5
0.7
0.9
Proportion
200 nodes
Time in log10(seconds)
1000
10
0.3
0.5
0.7
0.9
Proportion
300 nodes
Time in log10(seconds)
10000
100
0.3
0.5
0.7
0.9
Proportion
400 nodes
Time in log10(seconds)
1e+06
1e+04
1e+02
0.3
0.5
0.7
0.9
Proportion
500 nodes
Time in log10(seconds)
1000
100
10
0.5
0.7
0.9
Proportion
600 nodes
Time in log10(seconds)
10000
100
0.5
0.7
0.9
Proportion
700 nodes
Time in log10(seconds)
10000
1000
100
0.5
0.7
0.9
Proportion
800 nodes
Time in log10(seconds)
1e+05
1e+03
0.5
0.7
0.9
Proportion
900 nodes
Time in log10(seconds)
1e+05
1e+03
0.5
0.7
0.9
Proportion

**Figure S22. Distribution of the time (in log10) to execute the optimal algorithm in a family varying the number of nodes and the number of edges per node, signaled by the proportion of maximum edges per node.** The analysis with proportion 0.1 and 0.3 were removed due to the large computational time demanded in networks with a number of nodes bigger than 300 nodes.
